# Pharmacologically modified pluripotent stem cell-based cancer vaccines with anti-metastatic potential

**DOI:** 10.1101/2020.05.27.118471

**Authors:** Masae Heront-Kishi, Afag Asgarova, Christophe Desterke, Diana Chaker, Marie-Ghislaine de Goër de Herve, Ali G Turhan, Annelise Bennaceur-Griscelli, Frank Griscelli

## Abstract

Cancer is maintained by the activity of a rare population of self-renewing “cancer stem cells” (CSCs), which are resistant to conventional therapies. CSCs share several antigenic determinants with pluripotent stem cells (PSCs). We show here that PSCs, combined with a histone deacetylase inhibitor (HDACi), are able to elicit major anti-tumor responses in a model of highly aggressive breast cancer. This immunotherapy strategy was effective in preventing tumor establishment and efficiently targeted CSCs by inducing extensive modifications of the tumor microenvironment. The anti-tumor effect was correlated with a reduction in regulatory T and myeloid-derived suppressor cell populations and an increase in cytotoxic CD8^+^T cells within the tumor and the spleen along with a drastic reduction in metastatic dissemination and an improvement in the survival rate. These results demonstrate for the first time the possibility of using PSCs and HDACi as an allogeneic anticancer vaccine, in future universal immunotherapy strategies.

## INTRODUCTION

During the last decade, the concept of tumor heterogeneity has been extensively explored in solid tumors, leading to the identification, in several types of cancers, of a rare population of cells designated as “tumor-initiating cells” or “cancer stem cells” (CSCs) (Reya et al. 2001; Clarke et al. 2006; Al-Hajj et al. 2003). Within the bulk of a tumor, these cells represent a minor population with defined cellular and molecular characteristics: they are able to reinitiate tumor growth in immunodeficient mice, which suggests a capacity for self-renewal (Chen et al. 2012; Driessens et al. 2012) with a specific transcriptome (Puram et al. 2017; Galardi et al. 2016). Specifically, current evidence indicates that in addition to the well-known oncoprotein c-MYC, some of the key regulators of embryonic stem cells (ESCs), such as OCT4, SOX2, and NANOG, are also expressed in CSCs (Ben-Porath et al. 2008). These three factors participate in a highly integrated network along with c-MYC and polycomb proteins, that uses the epigenetic machinery to remodel chromatin through histone modification and DNA methylation. This ability to induce major epigenetic modifications was first demonstrated by groundbreaking experiments from the laboratories of Shinya Yamanaka and James Thomson; in these studies, researchers were able to induce embryonic features in somatic cells through the overexpression of a limited set of pluripotency genes, including *OCT4, SOX2, CMYC, KLF4, NANOG*, and *LIN28* (Takahashi et al. 2006; Yu et al. 2007). The induced pluripotent stem cells (iPSCs) obtained using this technology are closely linked to ESCs, as both express the same auto-regulatory circuitries (Robinton et al. 2012). Subsequent analyses of the genomic characteristics of iPSCs revealed that any genetic abnormalities present in later generations of cells could arise from the genomic status of the initial somatic target cell (Hussein et al. 2011; Schlaeger et al. 2015) or to their secondary appearance during their expansion *in vitro* (Hussein et al. 2011). In addition to the expression of pluripotency genes, this is another characteristic iPSCs share with CSCs, as random mutations in driver genes or outside of the functional genome have been shown to be present also in CSCs (Ben-Porath et al. 2008).

Because the antigen processing ability of CSCs is down-regulated, they express low levels of MHC-I molecules, which makes them difficult to detect by the host immune system (Prager et al 2019). This effect is further enhanced by the fact that the tumor microenvironment is highly immunosuppressive (Prager et al 2019). The combination of these two factors is remarkably effective in enabling CSCs to escape the surveillance of an efficient immune system.

In several cancers, gene expression programs have been identified that are similar to those of embryonic stem cells: namely, a “stemness” profile that is related to mesenchymal traits of carcinoma cells. These expression profiles include markers of the epithelial-mesenchymal transition (EMT) and low levels of MHC-I expression. Tumor cells undergoing EMT become CSCs with a capacity for early migration (and thus, metastasis) that are able to persist in a dormant stage for long periods of time. Such cancers correlate with aggressive, poorly differentiated tumor histology, invasive tumors, and very adverse outcomes (Ben-Porath et al. 2008; Glinsky et al. 2008; Schoenhals et al. 2009).

From these observations arose the idea of using ESCs or iPSCs as a source of tumor-associated antigens (TAAs), with the ultimate goal of provoking an anti-tumor immune response. The concept of using embryonic cells as a cancer vaccine is not new, but recent developments in ESC and iPSC technologies have facilitated work demonstrating their anti-tumor activity; these experiments have been primarily carried out in a preventive context using animal models of various cancers, including transplantable colon, lung, ovarian, and breast cancers (Li et al. 2009; Yaddanapudi et al. 2012; Zhang et al. 2013). More recently, autologous anti-tumor vaccines were developed using iPSCs, with TRL9 as adjuvant, in a prophylactic setting in a non-metastatic syngeneic murine breast cancer as well as in models of mesothelioma and melanoma (Kooreman et al. 2018). However, none of these existing reports evaluated the anti-metastatic potential of the anti-tumor vaccines, nor did they determine whether such vaccines have the ability to target CSCs and to modify their microenvironment. This question remains crucial, since anti-tumor vaccines that target only the bulk and transit-amplifying cells, but do not eradicate CSCs, are doomed to be unsuccessful. Here, using a model of aggressive triple-negative breast cancer, we show that pluripotent stem cells may be used as highly efficient whole-cell-based vaccines, in combination with a histone deacetylase inhibitor (HDACi), to prevent the establishment of CSC-enriched tumors and the development of lung metastases.

## RESULTS

### Poorly differentiated murine 4T1 breast tumors display an ESC-like expression signature *in vivo*

We first asked whether embryonic stem cell (ESC)-associated genes were enriched in the 4T1 murine breast cancer cell line. For this purpose, we generated a compendium of data that enabled us to compare different gene sets in different contexts, including adherent 4T1 cells cultured *in vitro* and 4T1 cells injected *in vivo* into the mammary fat pads of BALB/c mice. RNA from murine ESCs (D3) and from micro-dissected mammary glands were used as positive and negative controls, respectively.

From this transcriptome meta-analysis, we identified 1304 genes whose expression was significantly different among the four experimental groups (data not shown, differences determined using ANOVA). A comparison of implanted 4T1 cells and micro-dissected mammary glands highlighted 85 genes that were upregulated in the implanted 4T1 cells. By performing an unsupervised classification of the corrected matrix that contained all samples, we were able to identify two major clusters of genes: (i) those found in both non-tumoral mammary gland samples and 4T1 cells harvested in 2D culture *in vitro* and (ii) those found in both 4T1 cells implanted in BALB/c mice and in murine embryonic stem cells (mESCs) (**Figure 1A**).

**Figure 1.**
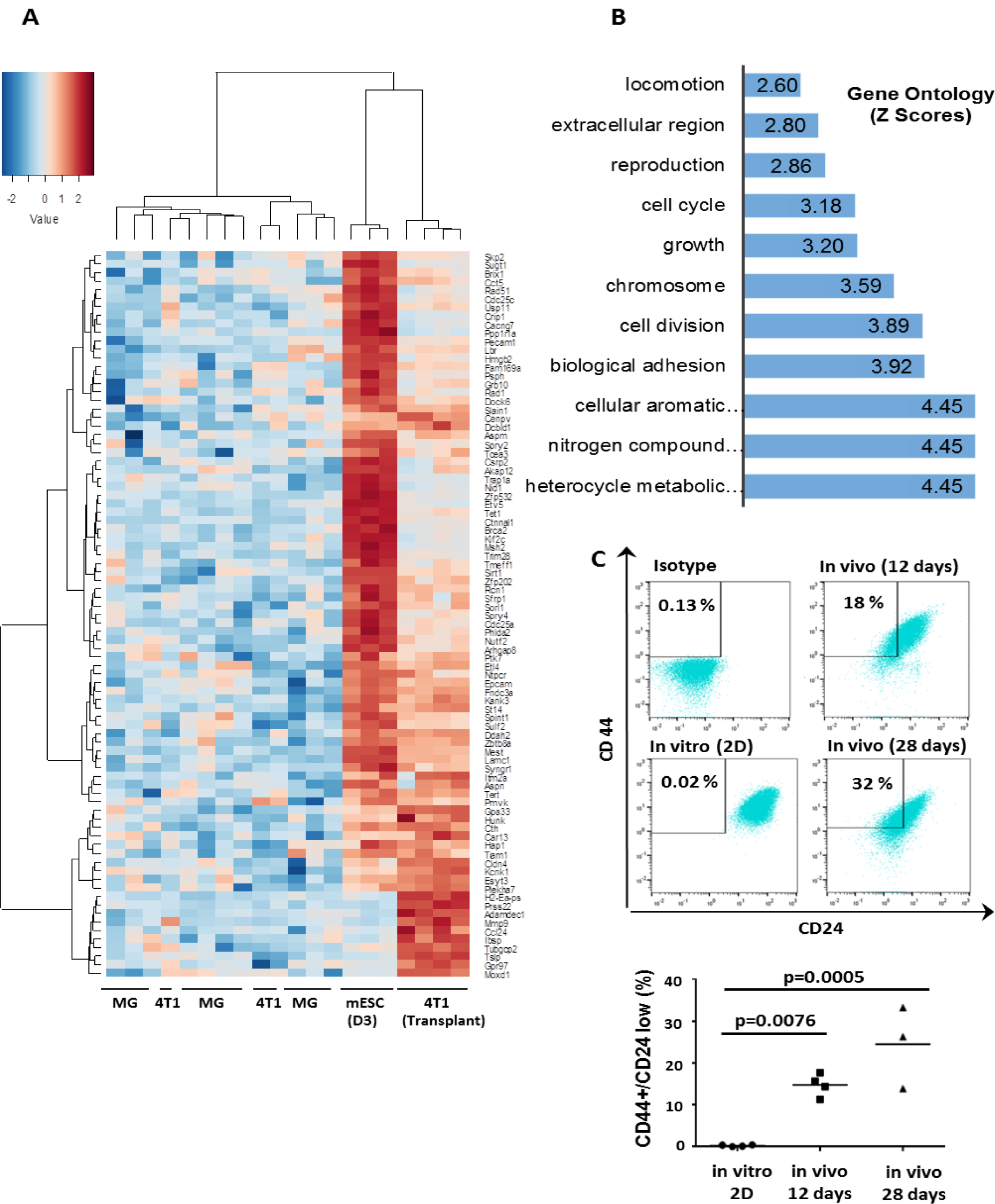
Breast tumors induced by 4T1 cells demonstrate an ESC-like expression signature *in vitro* and *in vivo*. (A) Transcriptome heatmap showing genes that were overexpressed in implanted 4T1 cells and in D3 murine embryonic stem cells (Euclidean distances, complete method). (B) Functional enrichment analysis of 85 genes that were overexpressed in implanted 4T1 cells and in D3 murine embryonic stem cells (Gene Ontology database); bars represent negative log10 of enrichment p-values. (C) Quantification of CD44/CD24 markers in 4T1 cells by flow cytometry *in vitro* and *in vivo* 12 and 28 days after implantation into the fad pat of BALB/c mice.

These results suggested that in the 4T1 cells, the transcriptional activation of embryonic genes that were shared with ESCs occurred only in the *in vivo* context, after the establishment of a tumor microenvironment. The functional enrichment of these stem-like cells was mainly linked to important processes in breast tumor development, such as aromatic metabolism, cell adhesion, cell growth, cell division, and cell cycle processes **(Figure 1B**). In order to confirm the stemness signature of implanted 4T1 cells, we used FACS analysis to quantify the expression of the breast cancer stem cell markers CD44 and CD24 (Al-Hajj et al. 2003) in 4T1 cells recovered *in vitro* and *in vivo* 12 and 28 days after injection into mammary fat pads. A CD44^+/high^/CD24^-/low^ population was identified in 4T1 cells *in vivo*, with a frequency up to 32% (**Figure 1C**), which was strong evidence of the emergence of a high proportion of CSC-like cells in this metastatic breast cancer model. The detection of shared antigens between 4T1 cells and ESCs prompted us to investigate whether ESCs could be used as tumor vaccines in this context.

### Vaccination with ESCs elicits anti-tumor activity associated with a tumor cell-specific CD8-dependent cytotoxic T lymphocyte response

To determine whether ESCs could stimulate the immune system to recognize embryonic antigens that are shared with tumor cells and thus confer protection against tumors, we used the aggressive triple-negative (ER^-^, PR^-^, HER2^-^) mouse 4T1 breast cancer model, which is known to have mesenchymal characteristics and to spontaneously and rapidly metastasize to the lungs (**Pulaski et al. 2001**). Naïve immunocompetent BALB/c mice were immunized two times (1 week apart) with irradiated human xenogeneic ESCs (H9) or murine allogeneic ESCs (D3); positive and negative control groups were injected with irradiated 4T1 cells and PBS, respectively. One week after the second injection, mice (n=35 mice) were challenged with 5×10^4^ 4T1 cells transplanted into the mammary fat pad (**Figure 2A**).

**Figure 2.**
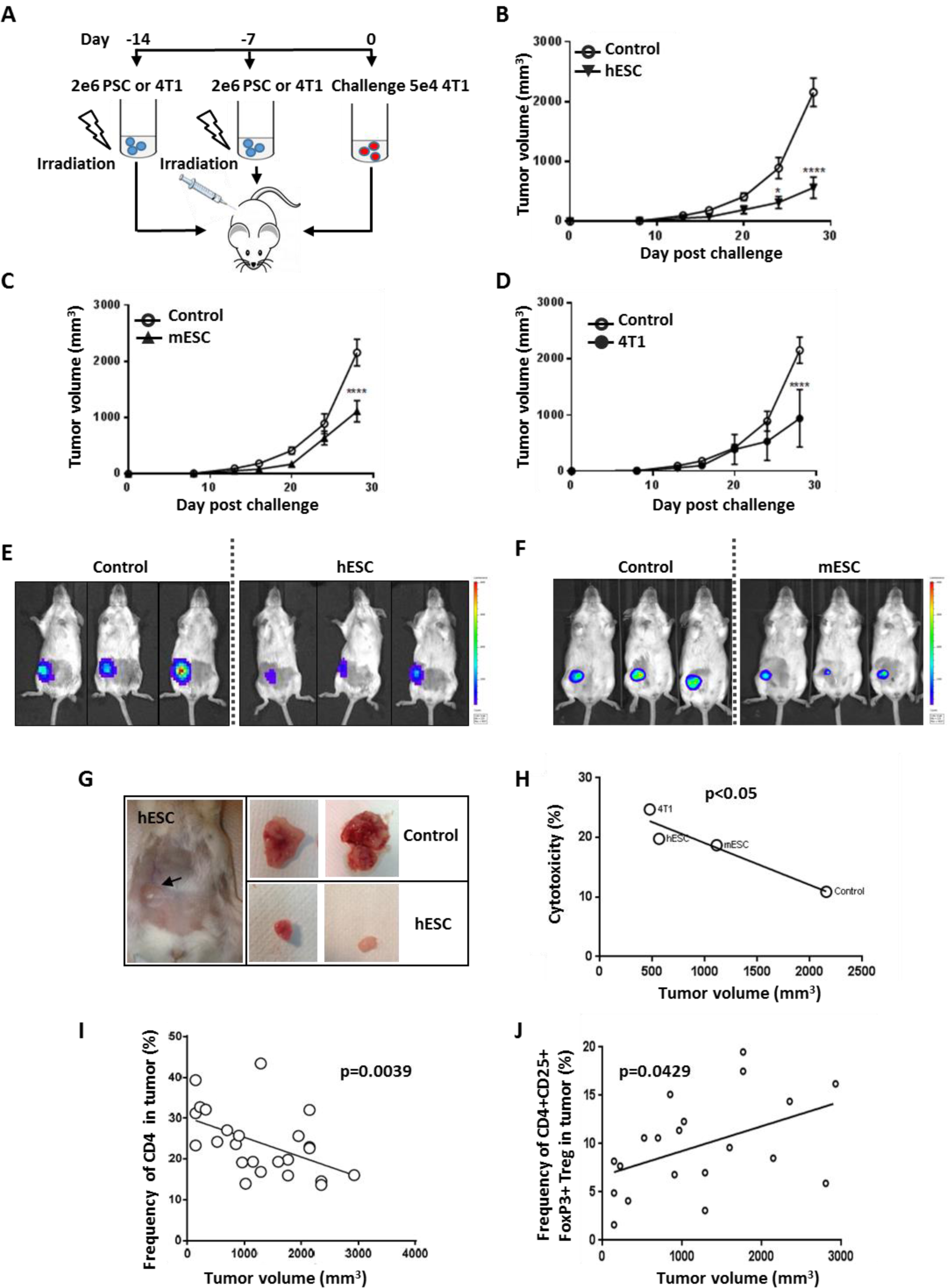
Vaccination with embryonic stem cells elicits anti-tumoral effect *in vivo*. (A) Vaccination and challenge protocol: irradiated stem cells or 4T1 cells were injected as a vaccine into BALB/c mice twice, with a one-week interval between doses. Mice were then challenged with 5×10^4^ 4T1 cells one week after the final dose. (B-D) hESC-, mESC-, and 4T1-vaccinated groups had significantly (p<0.0001) reduced tumor volumes compared to the control (PBS) group (n=5). (E, F) Bioluminescence imaging by IVIS of mice vaccinated with hESCs, mESCs. Macroscopic views of tumors from hESC-treated and untreated groups at 28 days post-challenge. hESC vaccination reduced tumor size compared to the control group. (G) A significant negative correlation was found between the frequency of cytotoxic CD8^+^ tumor-infiltrating lymphocytes and tumor volume (mm^3^) using the mean values of each treatment group (p< 0.05). (H) A significant negative correlation was found between the frequency of CD4^+^ tumor-infiltrating lymphocytes (%) and tumor volume (mm^3^). (I) A significant positive correlation was found between the frequency of infiltrated CD4^+^ CD25^+^ FoxP3^+^ Treg cells (%) in tumors and tumor volume (mm^3^).

Before conducting the experiment, we ensured that the ESCs that we used demonstrated the characteristics of pluripotent stem cells: they expressed pluripotency markers (**Figure S1A, S1B, S1C)** and formed teratomas *in vivo* that contained differentiated cells from the three germ layers (**Figure S1D**).

Mice were examined weekly, and their vital signs and tumor size was recorded. Twenty-eight days post-challenge, the tumors of mice that had been primed with hESCs, mESCs, and 4T1 cells were significantly (p<0.0001) smaller in volume compared to the tumors of non-immunized mice (**Figure 2B, 2C, 2D)**. The strongest effect was seen with xenogeneic hESCs (**Figure 2B, 2E, 2G**): tumors in this group were 73% smaller in volume than in the control group, compared to 52% smaller for allogenic ESCs (**Figure 2C, 2F**) and 56% smaller for 4T1 cells (**Figure 2D**). At this point, all mice were sacrificed so that we could analyze the phenotype of the immune cells present in the tumors and the spleens. This analysis revealed a significant negative correlation (p=0.05) between tumor volume and the amount of cytotoxic CD8^+^ T cells in the spleen (**Figure 2H**) as well as the amount of CD4^+^ T cell (p=0.0039) infiltrates in the tumor (**Figures 2I**). Moreover, a significant positive correlation was detected between tumor volume and the frequency of Tregs in CD4^+^ T cells in tumors (p=0.049) (**Figure 2J**). These results suggested that the anti-tumor effect observed with the use of stem cell-derived vaccines induced a cytotoxic T-cell response as well as a reduction in immune-repressive Treg cells in the tumor microenvironment. Our next step was to determine whether we could modulate the immune response induced by the vaccine by pharmacological priming or combination with an adjuvant.

### Treatment with valproic acid (VPA) enhances the immune response in 4T1 cells

We next propose to check the efficiency of the vaccine candidates *in vivo* in combination with HDACi such as VPA. To determine the potential effect of VPA on 4T1 cells, we first performed a transcriptome analysis on cells that had been treated with 0.5 mM of VPA for 10 days, and compared this to the transcriptome of untreated cells. These analyses identified 117 immune-related genes that were implicated in TNF alpha signaling and/or demonstrated a response to IFN-alpha and IFN-gamma **(Figure 3A).** These results were confirmed by a gene-set enrichment analysis that revealed significant enrichment in these three immune gene sets (**Figure 3B**). In addition, using the SAM algorithm we were able to identify 44 immune-related genes that demonstrated expression differences between the VPA-treated samples and their control counterparts (**Figure 3C, Table 1**). These were validated by principal component analysis (**Figure 3D**, p-value=3.3×10^−4^). Among these 44 immune genes, three had an expression fold-change greater than 2 such as *CD74, CCL2*, and *TNFRSF9* (**Figure 3E and Table 1**).

**Figure 3.**
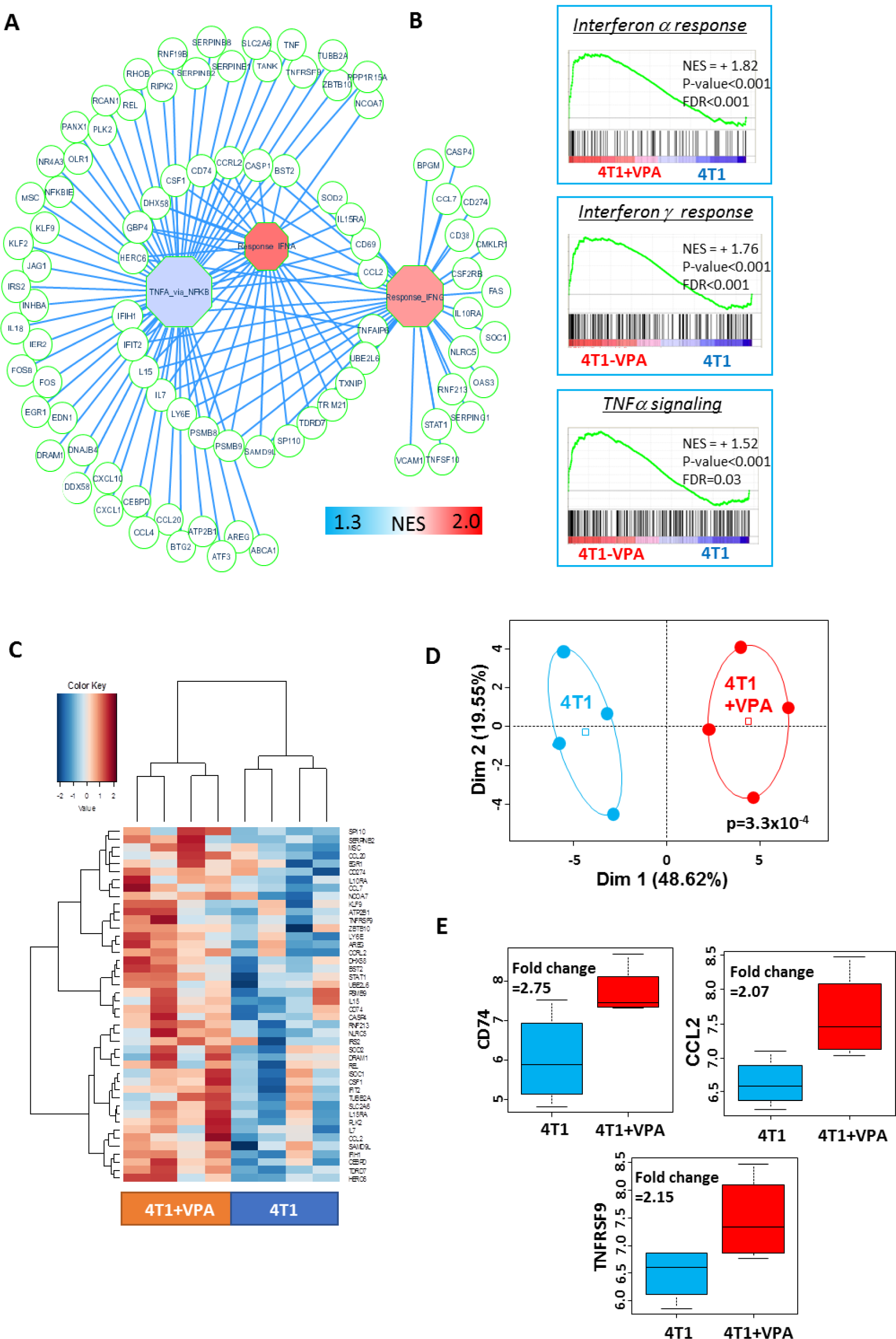
Treatment with valproic acid (VPA) induced an upregulation of the immune response in 4T1 cells *in vitro*. (A) A representation of the immune gene network upregulated by *in vitro* VPA treatment of 4T1 cells. Octagons represent enriched immune modules; circles are genes, which are connected to their respective enriched modules by blue edges (NES: normalized enrichment score). (B) Immune gene sets that were significantly enriched in VPA-treated 4T1 cells compared to untreated 4T1 cells (NES: normalized enrichment score, p-value and FDR were obtained by a hypergeometric test performed using the MSigDB 6.1 Hallmark database). (C) Expression heatmap depicting the immune genes that were significantly overexpressed in VPA-treated 4T1 cells (SAM algorithm, unsupervised classification on Euclidean distances with Ward method). (D) Unsupervised principal component analysis performed on upregulated immune genes in VPA-treated 4T1 cells. (E) Boxplot of three immune genes that were found to be significantly overexpressed as a result of VPA treatment (fold-change greater than 2).

More importantly, 4T1 cells that were treated with VPA expressed increased levels of MHC I in a dose-dependent manner (**Figure S2A**), which highlighted that VPA is able to enhance an anti-tumor immune response by improving recognition by T cells. In addition, measurements of relative fluorescence intensity with flow cytometry revealed that treatment with 2 mM of VPA resulted in a 2.1- and 2.7-fold increase in the expression of MHC I in 4T1 and mammosphere-derived 4T1 cells, respectively (**Figure S2B**). These results suggested that VPA treatment enhanced MHC I expression in both baseline and “stem-cell-like” contexts, and prompted us to investigate pluripotent cell vaccination in combination with VPA.

Before proceeding with the vaccination experiments, we asked whether treatment with only VPA had a significant effect on tumor size (**Figure S3A, S3B**) or on metastatic spread to the lungs (**Figure S3C**). We found that VPA alone had no significant anti-tumoral and anti-metastatic effects and did not change the levels of CD4^+^ or CD8^+^ T-cells in tumors, blood, or spleen. Similarly, levels of negative regulators of the immune system, such as Tregs and myeloid-derived suppressor cells (MDSCs), were unaffected by VPA treatment (**Figure S3D**). These data suggested that VPA alone was insufficient for inducing an immune response against the 4T1 model of breast cancer.

### ESC vaccination combined with VPA decreased lung metastases and led to recruitment of T cells

To determine whether a combination vaccine of hESCs+VPA provided better immunity against the epitopes shared between hESCs and 4T1 cancer cells, we evaluated the effectiveness of this approach compared to that using hESCs or VPA alone. In the two experimental groups (n= 24 mice) that were administered VPA, the drug was added in the drinking water at a dose of 4 mg/mL starting on the day of tumor challenge. Each mouse was injected with 5×10^4^ 4T1-GFP-Luc cells, which expressed the luciferase gene to enable quantification of viable tumor cells at metastatic sites.

In total, the experiment was conducted using 40 BALB/c mice, divided into four treatments: (1) PBS, (2) only VPA, (3) only hESCs (H9), and (4) hESCs+VPA. At day 44 after tumor implantation, the mice treated with only VPA showed a partial reduction (19.6%) in tumor volume compared to controls (treated with PBS) whereas the groups that received hESCs or hESCs+VPA had tumors that were significantly smaller (p<0.0001), with reductions compared to controls of 47.6% and 65.05%, respectively (**Figure 4A).** Bioluminescence imaging also confirmed that the groups treated with hESCs+VPA developed much smaller tumors compared to the untreated group (**Figure 4B**) and this effect was confirmed by analysis of the surgically removed tumors (**Figure 4C**). Since 4T1-GFP-Luc cells are able to rapidly give rise to lung metastases, we dissected the lungs (n = 24 mice) and quantified the total flux that originated from transplanted cells. We found that hESC vaccination combined with VPA treatment induced a highly significant reduction in pulmonary metastatic spread (more than 12-fold) compared to the control group (**Figure 4D, 4E**). In these experiments, tumor weight was found to be associated, in a highly significant manner (p=0.0062), with total flux in the lungs **(Figure 4F**). To evaluate immune cell infiltration in tumors, primary tumors and spleens were isolated and extensive immunophenotyping was performed. As shown in **Figure 4G**, tumors in mice that were pre-treated with hESCs+VPA contained significantly more CD4^+^ and CD8^+^ T-cells; the same result was found in the spleens of these animals (p=0.0179 and p=0.0186, respectively, for CD4^+^ T and CD8^+^ T cells) (**Figure 4H**). Furthermore, the expression of programmed cell death protein 1 (PD-1) in CD4^+^ and CD8^+^ T cells was significantly lower (p=0.012 and p=0.0171, respectively) in the spleens of hESC+VPA-treated mice (**Figure 4I, 4J**). Of all the treatments tested, the combined hESC+VPA regimen elicited the most anti-tumor activity, suggesting that this treatment successfully activated the immune system and caused significant modifications in the immunosuppressive microenvironment.

**Figure 4.**
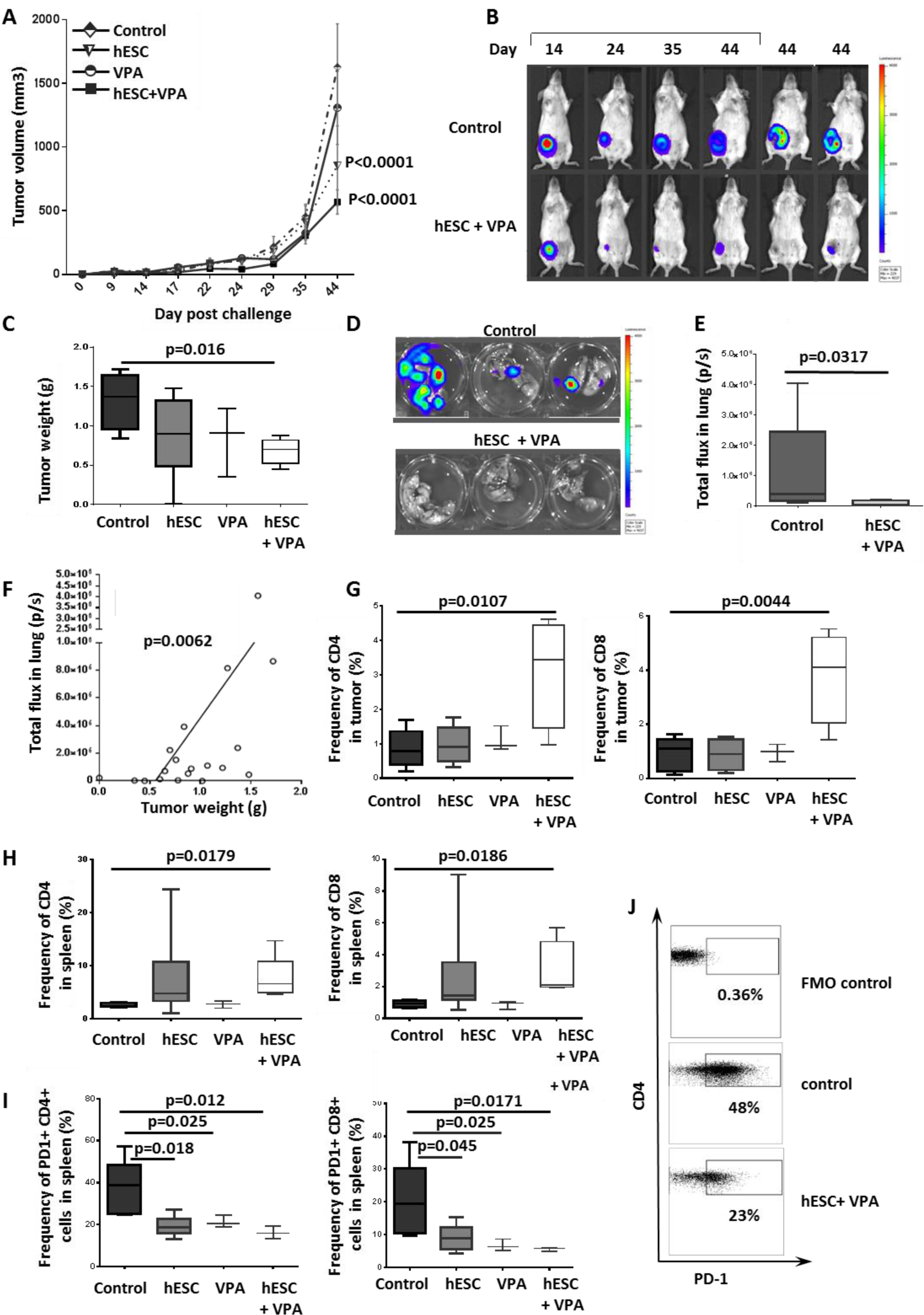
hESC vaccination with VPA administration had a synergistic effect on primary tumor growth and metastasis. (A) Tumor volumes in mice treated with VPA only, hESCs only, or hESCs+VPA (n=5-6). (B) Bioluminescence imaging by IVIS demonstrated the extent of the reduction in tumor areas in hESC+VPA-treated mice compared to the control group. On the left: the change over time in one mouse from each group. On the right: a comparison of tumors in different mice at day 44 post-challenge. (C) Comparison of tumor weights following surgical resection. (D) Direct bioluminescence imaging of surgically removed lungs (removed at day 44 post-challenge) revealed the reduction in metastastic load in mESC+VPA-treated mice compared to controls. (E) Bioluminescence imaging by IVIS revealed a significant decrease in lung metastasis in the hESC+VPA-treated group. (F) Measurements of total flux (photons/second; measured by IVIS) in surgically removed lungs had a significant positive correlation with tumor weight. (G) The frequency of CD4^+^ and CD8^+^ T cells was significantly higher in the tumors of hESC+VPA-treated mice than in other groups. (H) The frequency of CD4^+^ and CD8^+^ T cells in the spleen was significantly higher in the hESC+VPA-treated group compared to the control group. (I) PD-1 expression in CD4^+^ and CD8^+^ T cells was significantly lower in the hESC+VPA-treated group compared to the control group. (J) Dot plot images of flow cytometry results, showing high PD-1 expression by CD4^+^ T cells in the spleens of control mice, and reduced expression in the hESC+VPA group.

In a similar experiment, we also vaccinated mice with hESCs in combination with CpG ODN (**Figure S4A**), which is known to stimulate the Th1-based immune pathway by promoting the response of CD8^+^ T cells (Bode et al., 2011). However, there was no significant effect of treatment with hESCs+CpG ODN on tumor volume, tumor weight, or lung metastases compared to the control group (**Figures S4B-4F**).

### ESC vaccination combined with VPA decreased tumor size and lung metastases in the context of cancer stem cells

Next, we evaluated the cancer-protective effect of hESC vaccination combined with VPA administration by using a 4T1-derived mammosphere (MS) model to mimic CSCs. For this, we first produced MSs that contained a high proportion of CSC-like cells using two procedures; the only difference between the two was the addition (or not) of TGFβ and TNFα to the medium. After 5 days of culture, both protocols were effective in producing MSs (**Figure S5A)** that expressed aldehyde dehydrogenase 1 (ALDH1) (**Figure S5B**). As determined by flow cytometry, the amount of ALDH1 activity was 3.6- and 1.7-fold higher in MSs produced with or without TGFβ and TNFα, respectively, than in adherent 4T1 cells (**Figure S5C**). These experiments confirmed that treatment with both cytokines was effective in generating a CSC phenotype, as treated MSs demonstrated higher colony-formation ability and higher mammosphere-formation efficiency (**Figure S5D**), as well as higher expression of embryonic stem cell-associated genes such as *NANOG* and *SOX2* (**Figure S5E**). Likewise, MSs that were generated using TGFβ and TNFα exhibited higher tumorigenicity *in vivo*, which suggested that this cell population was enriched with more aggressive breast cancer stem/progenitor cells (**Figure S6A**).

We then applied this CSC model to our investigations of the tumor-protective effect of ESC vaccination combined with VPA. Mice were vaccinated with hESCs or mESCs and treated with VPA as described above, then challenged with MSs that had been produced from 4T1-GFP-Luc cells without the use of TGFβ and TNFα. On day 27 post-challenge, mice that had been treated with hESCs+VPA or mESCs+VPA had significantly smaller tumor volumes (p<0.0001 and p=0.0008 respectively) (**Figure 5A, 5B**) compared to the control group. In this model, xenogeneic hESCs were found to be more effective than allogeneic mESCs in preventing the establishment of 4T1-derived MSs. Bioluminescence imaging of tumors confirmed this result, as tumors in the hESC+VPA-treated group appeared much smaller than those in the control group (**Figure 5C**). Furthermore, mice treated with hESCs+VPA exhibited significantly (p<0.05) fewer lung metastases than the control group (**Figure 5D, 5E**). Immunological monitoring revealed a significant (p<0.05) increase in the frequency of CD4^+^ and CD8^+^ T cells (**Figure 5F, 5G)** and a significant decrease (p<0.05) in the abundance of Gr1^+^CD11b^+^ MDSCs in primary tumors of the hESCs+VPA treatment group compared to the control group (**Figure 5H and 5I**). The tumor specificity of cytotoxic T cells was further confirmed by the observation of increased secretion of interferon-gamma (IFN gamma) by tumor-infiltrating lymphocytes isolated from vaccinated mice after challenge with 4T1 MS cells. There was a significant negative correlation between IFN gamma secretion and tumor weight (**Figure 5J**), and mean induced IFNgamma values were significantly higher in the hESC+VPA-treated group than in either the control group or the mice primed with mESCs+VPA (**Figure 5K**).

**Figure 5.**
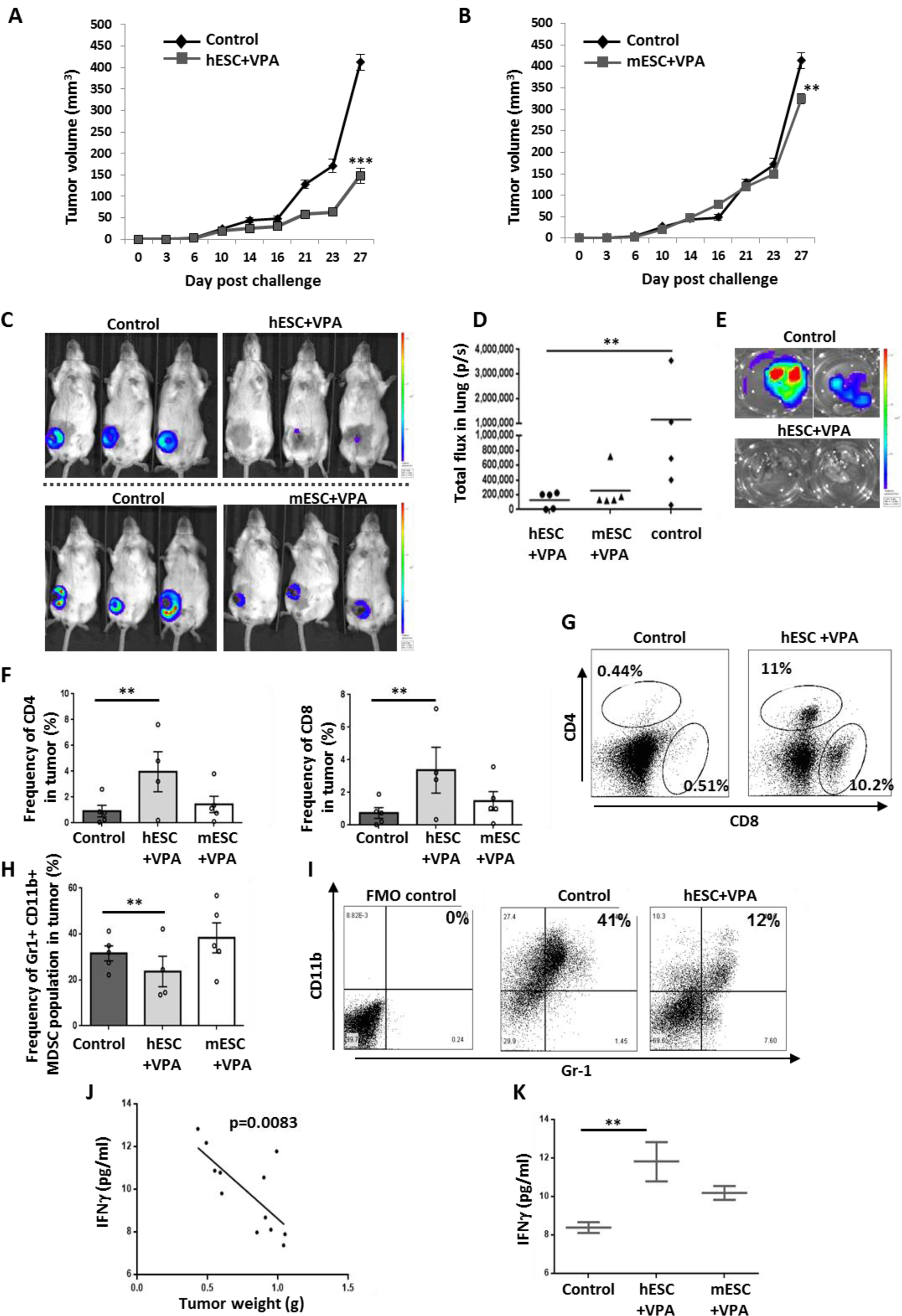
Evaluation of the anti-tumor and anti-metastatic potential of ESC+VPA treatment with respect to tumor stem cells. (A) hESC+VPA treatment reduced tumor volume (mm^3^) compared to controls at day 27 after challenge with mammospheres (n=5 per group). (B) mESC+VPA treatment reduced tumor volume (mm^3^) compared to controls at day 27 after challenge with mammospheres (n=5 per group). (C) Bioluminescence imaging of hESC+VPA and mESC+VPA treatment groups compared to control group. (D) Lung metastases in treated (hESCs+VPA, mESCs+VPA) and control mice, as shown by total flux (photons/second) (E) Bioluminescence images of the surgically removed lungs of control and hESC+VPA-treated mice. (F) Quantification by flow cytometry of CD4^+^ and CD8^+^ tumor-infiltrating lymphocytes from treated and untreated mice. (G) Dot plot images of flow cytometry results, showing an increase in CD4^+^ and CD8^+^ T cells in mice treated with hESC+VPA compared to the control group. (H) Quantification of Gr1^+^CD11b^+^ cell populations in tumors of mice treated with ESCs+VPA compared to the control group. (I) Dot plot images of flow cytometry results, showing the double Gr1^+^CD11b^+^ cell populations in tumors in control mice and in mice treated with hESC+VPA. (J) A negative correlation was found between the secretion of IFN-γ (pg/mL) by tumor-infiltrating lymphocytes and tumor burden. (K) Quantification of IFN-gamma (pg/mL) secretion by tumor-infiltrating lymphocytes in treated and control mice.

We repeated the experiment using MSs generated with TGFβ and TNFα, and obtained similar results: drastic reductions in tumor development following vaccination with either hESCs+VPA or mESCs+VPA (**Figure S6B and S6C**). With this model we also observed a significant reduction of cancer cell dissemination in lungs (**Figure S6D, S6E, S6G**).

### Anti-tumor effects of iPSCs in an autologous or allogeneic context

We next wished to evaluate the anti-tumor activity elicited by the same protocol, but with iPSCs instead of ESCs. For this purpose, we generated murine iPSCs (miPSCs) from BALB/c fibroblasts using lentiviral vectors that expressed OCT4, KLF4, SOX2, and c-MYC. We obtained fully reprogrammed miPSCs that exhibited iPSC characteristics with respect to morphology (**Figure S7A**), gene expression, and surface markers. Specifically, these cells strongly expressed OCT4, and NANOG (**Figure S7B**) and SSEA1 (**Figure S7C**), and were able to generate teratomas *in vivo* (**Figure S7D**).

We then used these cells to conduct the same vaccination experiment as performed with hESCs (**Figure 2A**), but also investigated whether the results differed if mice received autologous or allogeneic miPSCs. At day 20, mice treated with autologous miPSCs, but not VPA, had significantly smaller (35%; p=0.0058) tumors compared to controls, whereas mice vaccinated with allogeneic miPSCs were less strongly affected (reduction of 21%) (**Figure 6A and 6B**). Instead, mice vaccinated with either autologous or allogeneic miPSCs in combination with VPA treatment demonstrated a drastic reduction in their tumor burdens, of 48% (p=0.0018) and 61% (p<0.0001), respectively, compared to control mice (**Figure 6A and 6B**).

**Figure 6.**
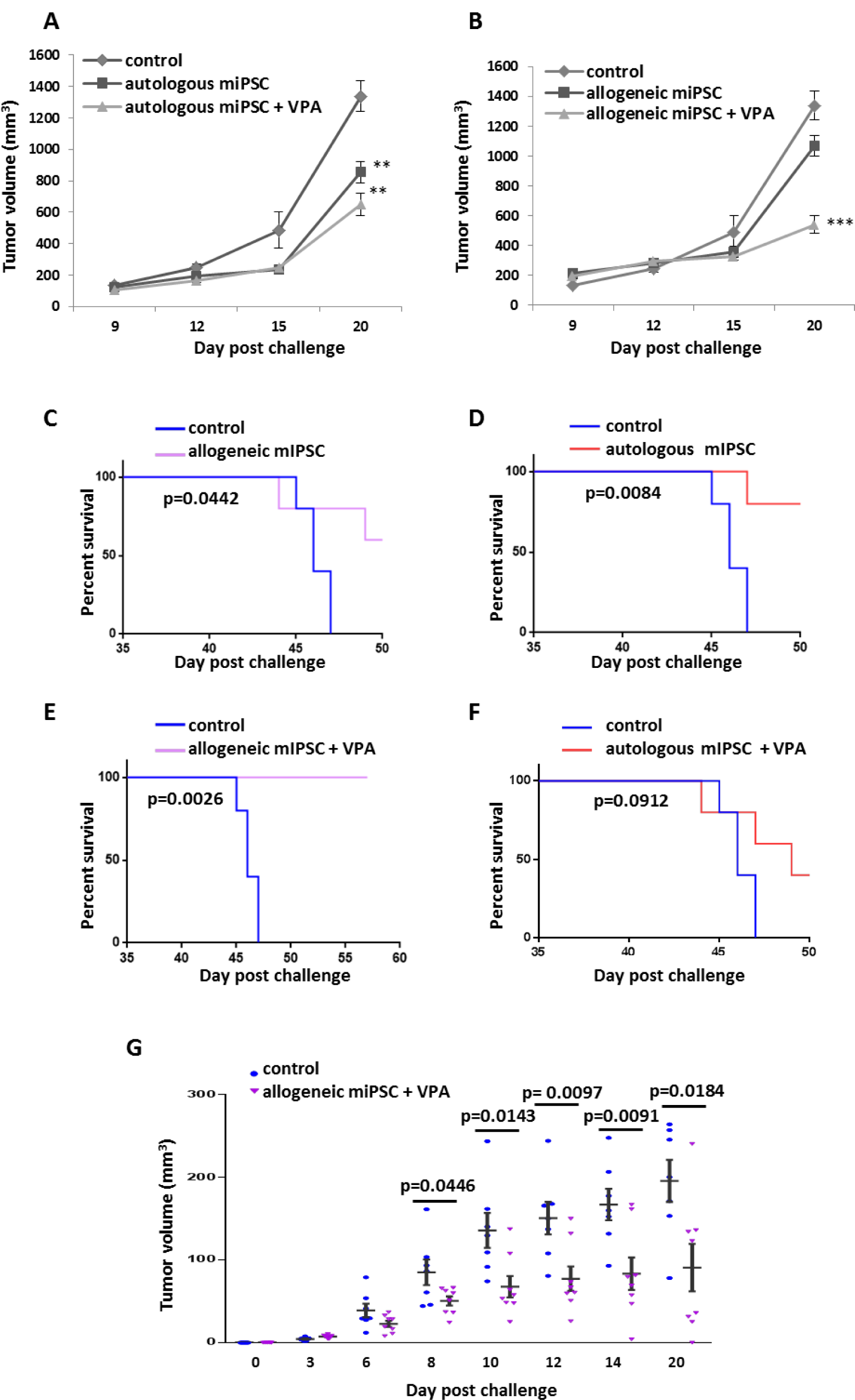
Evaluation of the anti-tumor effects of murine iPSCs derived from BALB/c or C57Bl/6 mice. (A) Tumor growth in mice that were immunized with murine BALB/c-derived iPSCs, with or without VPA, compared to control group. (B) Tumor growth in mice that were immunized with murine C57BL/6-derived iPSCs, with or without VPA, compared to control group. (C) Effect of allogeneic miPSC treatment on the survival of mice challenged with 4T1 cells. (D) Effect of autologous miPSC treatment on the survival of mice challenged with 4T1 cells. (E) Effect of allogeneic miPSC+VPA treatment on the survival of mice challenged with 4T1 cells. (F) Effect of autologous miPSC+VPA treatment on the survival of mice challenged with 4T1 cells. (G) Tumor volumes of mice treated with allogeneic miPSCs+VPA compared to those of untreated mice. The data represent the mean ± SEM of tumor volumes (8 mice per group).

In this experiment, we also performed a long-term survival study. Vaccination with either type of miPSC conferred a significant improvement in survival rate over the control group (**Figure 6C and 6D).** In addition, a significant survival benefit was observed when VPA was combined with the allogenic vaccine (**Figure 6E**), but this effect was less pronounced for autologous miPSCs+VPA (**Figure 6F**).

To investigate this further, we performed additional experiments in which we vaccinated mice (n = 8) with 2×10^6^ C57BL/6-derived allogenic miPSCs and compared tumor progression with that of unvaccinated mice (n = 8). All mice were challenged with 5×10^4^ 4T1 cancer cells using the initial protocol (**Figure 2A**), and tumor size was monitored until day +20 (**Figure 6G**). After 3 days, all mice from both groups presented tumors of similar size. In 7 of the 8 control mice, tumors grew dramatically over the remainder of the experiment. Instead, in mice that were treated with the allogeneic miPSC vaccine, tumors grew much more slowly, and indeed, even shrank in 4 of the 8 treated mice (**Figure 6G**). In one mouse, the tumor completely disappeared, while the other three mice demonstrated a partial regression in tumor size, with final volumes of less than 35 mm^3^ (**Figure 6G**).

### Generation of an efficient memory immune response after vaccination with miPSCs

To investigate whether vaccination with miPSCs could induce a T cell memory, we inoculated mice with 2×10^6^ irradiated BALB/c-derived miPSCs, a total of 6 times, once every 30 days. Thirty days (**Figure S8A**) or 120 days (**Figure 7A**) after the final inoculation, mice were challenged with 5×10^4^ 4T1-GFP-Luc cells. All vaccinated mice received 4 mg/mL VPA by oral route starting from the day of tumor injection, and tumor growth was monitored for 26 or 28 days after challenge. In the mice challenged with 4T1 cancer cells 30 days after the final vaccine dose, tumor volume was reduced by 64% compared to unvaccinated mice (695±102 mm^3^ versus 1968±96 mm^3^, p=0.005; **Figure S8B**). When we increased the time elapsed between the final vaccine dose and tumor implantation to 120 days, we observed again a significant response (46% reduction: 320±23 mm^3^ versus 600±40 mm^3^ for controls, p<0.0001; **Figure 7B**). In both cases, vaccination with miPSCs was able to generate an effective immune response that resulted in a significant inhibition of tumor growth compared to the control group. To better understand the mechanisms underlying this effect, we performed an in-depth investigation of the mice for which 120 days elapsed between their final vaccine dose and tumor challenge. An analysis of metastatic dissemination revealed a major reduction in the extent of lung metastases in this group compared to controls (**Figure 7C**), with a significant correlation between tumor burden and metastatic spread (**Figure 7D**). Treatment with miPSCs+VPA was also correlated with a significant increase in the frequency of CD4^+^ and CD8^+^ T cells in tumors (**Figure 7E)** and in spleens (**Figures S9A, S9B**). Furthermore, we observed a significant decrease in the frequency of PD1^+^ CD4^+^ cells in the spleen (**Figure S9C**), as well as a decrease in PDL1 expression in tumor cells (**Figure 7F**), which was strong evidence that our vaccination protocol had a negative impact on T-lymphocyte anergy. Similarly, miPSC+VPA treatment led to a decrease in the frequency of CD4+ CD25+FoxP3+ Tregs (**Figure 7G)** and in different populations of Arg1^+^ CD11b+ Gr1+ preMDSCs and Arg1^+^ CD11b+ Ly6+ gramMDSCs (**Figure 7H**), which indicated that vaccination had a negative impact on the main immunosuppressive actors of the immune system.

**Figure 7.**
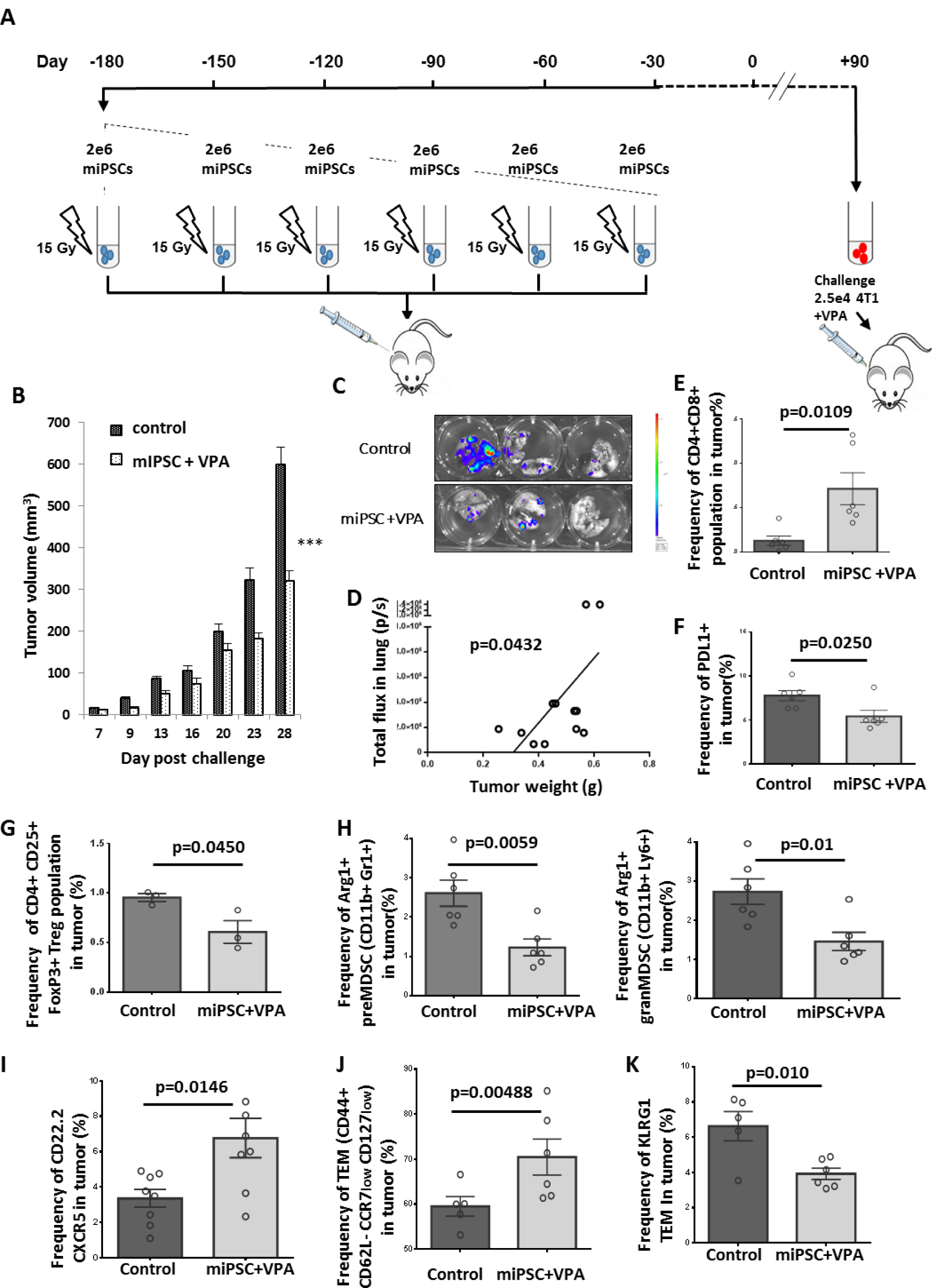
Effective memory immune response following vaccination with miPSCs. (A) Experimental protocol to evaluate *in vivo* immune memory generated by vaccination: BALB/c mice were injected subcutaneously six times with 2×10^6^ miPSCs (15Gy irradiated) in the right flank. Tumor challenge occurred 120 days after the final vaccine dose was given; mice were injected with 2.5×10^4^ 4T1-GFP-Luc cells and VPA was added in their drinking water. (B) At day 28 post-challenge, breast tumors were significantly smaller in mice that had undergone the six-month vaccination protocol compared to unvaccinated mice (n=6 per group). (C) Bioluminescence images of lungs isolated from miPSC-vaccinated and control mice 28 days after tumor challenge. (D) A significant correlation was found between tumor burden and metastatic spread in the lungs of vaccinated mice at day 28 post-challenge. (E) The frequency of CD4^+^CD8^+^ cells in tumors of treated and untreated mice, as measured by flow cytometry. (F) The frequency of PDL1^+^ cells in tumors of miPSC-vaccinated mice compared to controls, as measured by flow cytometry. (G) The frequency of Treg cells in tumors of miPSC-vaccinated mice compared to controls, as measured by flow cytometry. (H) The frequency of Arg1^+^ preMDSCs and granMDSCs in tumors of miPSC-vaccinated mice compared to controls, as measured by flow cytometry. (I) The frequency of CXCR5^+^ CD22.2^+^ LB cells in tumors of miPSC-vaccinated mice compared to controls, as measured by flow cytometry. (J) The frequency of T-effector memory cells in tumors of miPSC-vaccinated mice compared to controls, as measured by flow cytometry. (K) The frequency of KLRG1^+^ T-effector memory cells in tumors of miPSC-vaccinated mice compared to controls, as measured by flow cytometry.

Among the other long-term effects of the miPSC+VPA treatment, we also detected potential effects on CD22.2^+^ B-lymphocytes. Specifically, we observed a significant increase in CXCR5^+^ B cells in vaccinated mice compared to controls (**Figure 7I**), which suggested that B-lymphocytes had migrated to tumor sites as a result of our treatment. These observations prompted us to evaluate changes in the levels of T-effector memory cells (TEMs). We found that miPSC-VPA treatment led to an increase in the frequency of CD44+ CD62L-CCR7low CD127low TEMs in tumors (**Figure 7J**). We also observed a decrease of the expression of PD1 by TEMs (CD44+CD62L-CCR7 low CD127 low) from spleens (**Figure S9D**), as well as in KLRG1 expression by TEMs from tumors (**Figure 7K**), which suggested that the immune memory had reverted to an active state, decreasing both senescence and anergy. These results provide strong evidence that injection with a miPSC vaccine led to the establishment of long-term immune memory and the activation of effective anti-tumor immunity. This benefit could also be seen in survival data, as our vaccination protocol dramatically improved the long-term survival of treated mice compared to controls (median survival was 54 days in control group but not reached in vaccinated group) (**Figure S8C).**

Finally, we wished to consolidate all of these results by carrying out the same vaccination protocol and then challenging mice 120 days after the final vaccine dose with 5×10^5^ MS-derived 4T1 cells (**Figure S10A**). As before, 4 mg/mL VPA was added in the drinking water starting on the day of tumor injection. As we found with our previous results, immunization with miPSCs led to a significant (p<0.0001) and highly effective immune response to MS-derived 4T1 cancer cells. The total tumor burden in treated mice was reduced by 83% compared to controls, with a massive reduction in tumor sizes (**Figure S10B, S10C**). The long-term immune memory generated by our protocol also significantly protected the mice from developing lung metastases (**Figure S10D**); as before, we observed a significant correlation between tumor weight and metastatic dissemination (**Figure S10E**).

To investigate the underlying mechanisms of the long-term immune protection that resulted from the vaccination protocol, we performed a transcriptome analysis of tumors from untreated mice and from mice primed with 6 doses of miPSCs and VPA in order to find genes that were differentially expressed between the two conditions. As can be seen in **Table 2** and **Figure 8A**, 206 genes were found to be differentially expressed in a highly significant manner (**Table 2, Figure 8A**). Of the genes that were upregulated in treated mice, 98 were implicated in the immune response to cytokines and lymphocyte chemotaxis (**Figure 8B**). When we analyzed genes linked to the immune response, we detected a significant enrichment in monocyte-derived dendritic cells (Normalized Enriched Score (NES) =1.66, p-value<0.001), T cells (NES=1.54, p-value<0.001), and B cells (NES=1.33, p-value<0.001) (**Figure 8C**). Likewise, a gene network-based analysis supported the major contribution of B-cells and T-cells by highlighting genes associated with these respective infiltrations (**Figure 8D**). Interestingly, a significant upregulation (8.8-fold; p-value=5.1787803E-5) of chemokine (C-X-C motif) ligand 13 was detected in tumors that persisted after vaccination (**Table 2**). This result was confirmed by qRT PCR, which also revealed a strong upregulation of CXCL13, CXCL9, and CXCL10 in the tumors of vaccinated mice (**Figure S11**). Together, these data strongly suggest that the essential mode of action of the vaccine is to recruit B and T lymphocytes to tumor sites via an active immune response that utilizes CXCL9, CXCL10 and CXCL13.

**Figure 8.**
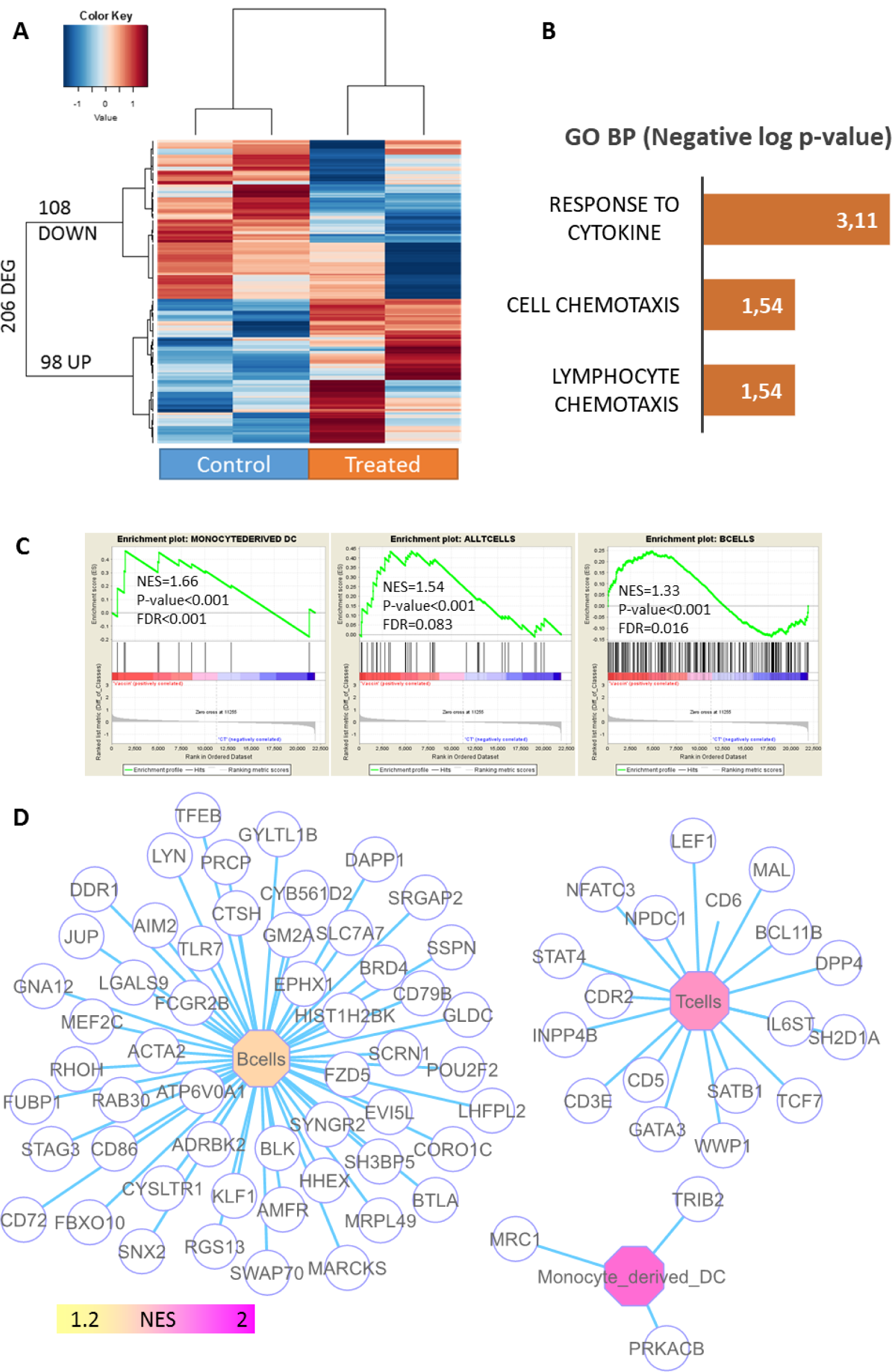
Profiling of immune-associated genes in 4T1 tumors in mice primed with miPSCs. (A) Differentially expressed genes (DEGs) between control and vaccinated mice xeno-transplanted with 4T1 cancer cells; expression heatmap was produced with an unsupervised classification algorithm (Euclidean distances, Ward method). (B) Barplot of functional enrichment in Gene Ontology Biological Processes of genes that were overexpressed in vaccinated mice. (C) Immune profiling carried out by gene-set enrichment analysis on the transcriptome of 4T1-transplanted mice (NES: normalized enriched score) (D) Genes associated with the immune network that were enriched in 4T1 tumors from miPSC+VPA-treated mice.

## MATERIALS AND METHODS

### Cell line isolation and maintenance

An hESC line (H9) was obtained from the University of Wisconsin (Thomson et al. 1998) and used under agreement #RE07-008R with the French Biomedical Agency. These cells were maintained on mitomycin-treated mouse embryonic fibroblasts (MEFs) in Dulbecco’s modified Eagle’s/F12 medium (DMEM) that was supplemented with 20% Knockout serum replacement, 1% penicillin and streptomycin (Invitrogen), 1% non-essential amino acids, 1 mM 2-mercaptoethanol (Sigma), and 12.5 µg/mL of basic fibroblast growth factor (bFGF). A murine iPSC line was generated from fibroblasts isolated from primary cells of BALB/c mice via transduction with Cre-Excisable Constitutive Polycistronic Lentivirus, which expresses OCT4, SOX2, CMYC, and KLF4 (EF1alpha-STEMCCA-LoxP backbone from Millipore). An iPS cell line from C57BL/6 mice was purchased from ALSTEM (Richmond, CA). Murine ESCs (D3) and murine iPSC lines were maintained on mitomycin-treated MEFs in DMEM glutamax (Gibco) that contained 15% fetal bovine serum (Eurobio), 1% penicillin and streptomycin (Invitrogen), 1 mM 2-mercaptoethanol (Sigma), and 1000 units/mL leukemia inhibitory factor. Prior to vaccine preparation, hESCs were cultured on Geltrex (Thermo Fisher Scientific) in Essential 8 medium (Thermo Fisher Scientific), while D3 cells and murine iPSCs were cultured on gelatin (Sigma) in the same medium. For the teratoma assays, iPSCs were resuspended in PBS and injected at a dose of 2×10^6^ cells per mouse to the flank of immunodeficient mice (NOD-SCID). Mice were sacrificed two months later for histological analysis of teratomas. All PSCs and 4T1 cells used for vaccination were irradiated with a lethal dose of 15 Gy and 75 Gy, respectively, prior to injection.

The breast cancer line 4T1 was obtained from ATCC (CRL-2539) and grown in DMEM with 10% FBS and 1% penicillin and streptomycin under normal culture conditions. For *in vitro* induction of mammospheres (MSs), the 4T1 cells were seeded in low-attachment 6-well plates at a density of 100,000 cells per well with a cocktail of MEF-conditioned medium (3/4 MEF-conditioned medium + 1/4 mES medium + 4 ng/mL bFGF), with or without the addition of the cytokines TNF-alpha (20 ng/mL, Cell Signaling Technology) and TGF-beta 1 (10 ng/mL, Cell Signaling Technology). The 4T1 cell line was transduced using the retroviral vector pMEGIX, which encodes the genes for green fluorescent protein (GFP) gene and luciferase. Stable clones were isolated and selected by GFP expression using flow cytometry.

### Transcriptome meta-analysis of 4T1 cells

In order to evaluate the stem-cell signature of the 4T1 murine breast cancer cell line, we compared the transcriptomes of 4T1 cells in different contexts to that of the murine embryonic stem cell line (D3, from GEO dataset GSE51782) and of micro-dissected samples of mammary glands (from GEO dataset GSE14202) (Padovani et al. 2009). Microarray data of *in vitro*-generated and murine-transplanted 4T1 samples were obtained from GEO accessions GSE73296 and GSE69006, respectively. Each normalized matrix was merged with its respective annotated matrix via their gene symbol annotation, and processed for batch correction using the Stanford algorithm in R (Tibshirani et al. 2002). One-way analysis of variance (ANOVA) with 500 permutations was performed on the four experimental groups using the corrected matrix. A nested analysis of differentially expressed genes (DEGs) was performed on 4T1-transplanted and mammary-gland samples using a significance analysis of microarrays (SAM) with a maximum false discovery rate (FDR) threshold of 5 percent (Tusher et al. 2001). Subsequently, functional enrichment was identified using the Gene Ontology Database with the Go-elite standalone application (Zambon et al. 2012). A DEG heatmap was created using the made4 Bioconductor package (Culhane et al. 2005) in R version 3.4.1.

### Transcriptome analysis of 4T1 cells treated with valproic acid

Total RNA was extracted following the instructions of the manufacturer (TRIzol, Life Technologies) from 4T1 cells cultured with and without valproic acid (VPA) at a dose of 0.5 mM for 10 days. The concentration of total RNA was measured on a Nanodrop device and the quality of the extracted nucleic acid was assessed using a Bio-analyzer 2100 (Agilent technologies, CA). Microarray probes were synthetized in one cycle of RNA amplification in which molecules were labeled (Affymetrix microarray station, Affymetrix, CA). The labeled microarray probes were hybridized on a Mouse Clariom S (mm10) microarray (Thermo Fisher Scientific), and the CEL files of microarray data obtained from the Affymetrix platform were normalized using the RMA method in Affymetrix Expression Console software (Affymetrix, CA). Gene-set enrichment analysis was performed with the online java module of GSEA software, version 3.0, while a network-based gene-set enrichment analysis was performed with Cytoscape software, version 3.6.0. Bioinformatics analyses were performed in R version 3.4.1; the R-package made4 was used to create an expression heatmap using Euclidean distances and the Ward method, and the FactoMineR R-package was used to perform an unsupervised principal component analysis. Genes with significant differences in expression were selected with the SAM algorithm using a FDR threshold of 5 percent.

### Animal model

Wild-type female BALB/c mice, 8-10 weeks old, were purchased from Janvier Laboratory and maintained at the animal core facility using standard guidelines. All protocols for animal experiments were approved by the Animal Care Committee of the Val de Marne. *In vivo* tumor studies were performed using 5 to 10 BALB/c mice in each group unless otherwise specified. Mice were anesthetized using 2% isoflurane in 100% oxygen and injected subcutaneously with 2×10^6^ irradiated ESCs or iPSCs (suspended in 100 µl of PBS) in the right flank; each mouse was vaccinated twice, with a one-week interval between injections. One week following the second vaccine dose, 5×10^4^ 4T1-GFP-Luc cancer cells were resuspended in 100 µL PBS and injected into mice at the 4th mammary fat-pad using a tuberculin syringe. For the H9+CpG vaccine, irradiated H9 cells were suspended in 100 µL PBS that also contained 50 µg CpG (Invivogen). After tumor implantation, VPA was orally administered to vaccinated mice through their drinking water at a dose of 4 mg/mL. Specifically, VPA (Sigma) was dissolved in drinking water at a concentration of 0.4% w/v and provided to mice *ad libitum* in feeding bottles, an approach that leads to approximate concentrations of 0.4 mM VPA in plasma, as previously reported (Shabbeer et al., 2007). Tumor growth was measured using a caliper. *In vivo* bioluminescence imaging was performed using an IVIS Spectrum system (Perkin Elmer) and images were analyzed and quantified with Living Image software (Perkin Elmer). For direct imaging of lungs at sacrifice, lungs were isolated and incubated with 150 µg/mL luciferin in 12-well plates and then imaged. After sacrifice, tumors were dissociated using a dissociation machine (GentleMACS dissociator, Miltenyi Biotec) and the Mouse Tumor dissociation kit (Miltenyi), without enzyme R (Tumor dissociation kit, Miltenyi Biotec). Spleens were dissociated using a cell strainer (Fisher), after which RBC Lysis Buffer (eBioscience) was used to remove red blood cells. After dissociation, the cell suspension was washed with PBS and used in subsequent analyses.

To study the ability of the vaccine product to induce immunological memory, BALB/c mice were immunized subcutaneously with 2×10^6^ irradiated miPSCs in 100 µl PBS solution six times (3-day intervals between administrations); the control group received only PBS. Thirty or 120 days following the final vaccine dose, mice from both groups were injected with 2.5×10^4^ 4T1-GFP-Luc cancer cells in the 4th mammary fat-pad. Following tumor implantation, vaccinated mice were also treated with VPA through their drinking water (4 mg/mL, Sigma).

### Staining of immune cells and tumor cells for FACS analysis and ELISA assay

Cells that had been isolated from spleen and tumor tissues were resuspended in PBS (Gibco) that contained 1% FBS. They were then stained for cell surface markers using antibodies for CD44, CD24, CD45, CD8a, CD25, CD279 (PD-1), MHC class I (eBioscience), CD11b, Gr-1, CD3, CD4, CD279 (PD-1), CD45, CD44, and CD24 (Miltenyi Biotec). Dead cells were excluded using 7-aminoactinomycin D (7-AAD, eBioscience) and zombie violet (BioLegend). FACS analysis was conducted using a MACSQuant analyzer (Miltenyi Biotec). Immune stimulation was performed using 50 ng/mL Phorbol 12-Myristate 13-Acetate (PMA) (Sigma) and 500 ng/mL ionomycin b (Sigma) in RPMI 1640 medium (Gibco) that contained 10% bovine fetal serum (Gibco) and 1% penicillin and streptomycin. IFN-gamma was quantified using the IFN-gamma kit (eBioscience) following the manufacturer’s protocol.

### Aldefluor assay

The Aldefluor kit (Stem Cell Technologies) was used to characterize the ALDH activity of 4T1 cells as described in (Pearce et al. 2005). Cells were incubated at 37°C in Aldefluor assay buffer, which contained the ALDH substrate BODIPY-aminoacetaldehyde. To determine baseline fluorescence, the enzymatic activity of ALDH was blocked by the inhibitor DEAB, and these data were used in determining the gates for FACS analysis. FACS analysis was performed using a MACSQuant analyzer (Miltenyi Biotec).

### Cytotoxicity assays

4T1 target cells were incubated for 9 min with 2 µM Carboxyfluorescein succinimidyl ester (CFSE) (eBioscience) in PBS. Positive selection was performed to isolate CD8^+^ T cells from primary tumors and spleens using CD8a microbeads (Miltenyi Biotec), an LS column, and a MACS separator (Miltenyi Biotec), according to the manufacturer’s protocol. In total, 10,000 CFSE-labeled 4T1 cells and 10,000 CD8^+^ sorted cells from primary tumors or spleens were plated with RPMI, 10% FBS, and 1% penicillin and streptomycin (Invitrogen) in 98-well plates, then incubated at 37°C overnight. The following day, the incubated cells were stained with 7-AAD to assess viability, then subjected to flow cytometry analysis to determine the fraction of live, CFSE-positive cells. The cytotoxicity rate (%) was calculated using the T/C ratio = % target cells / % control cells (non-immunized cells), survival rate (%) by the formula = {T/C ratio of sample}/{T/C ratio of untreated BALB/c mouse}, and cytotoxicity rate (%) by the formula = 100 – survival rate.

### Transcriptome analysis of 4T1-derived tumors from mice vaccinated with miPSCs and VPA

From the 4T1 tumors that had been surgically removed from mice in the experimental (vaccinated with miPSCs and treated with VPA) and control groups, we extracted total RNA following the instructions of the manufacturer (TRIzol, Life Technologies). Concentration of total RNA was measured using a Nanodrop spectrophotometer and the quality of the extracted nucleic acid was assessed on a Bio-analyzer 2100 (Agilent Technologies, CA). Microarray probes was synthetized by one cycle of RNA amplification in which molecules were labeled in an Affymetrix microarray station (Affymetrix, CA). Labeled microarray probes were hybridized on a Mouse Clariom S (mm10) microarray (Thermo Fisher Scientific). The CEL files of microarray data obtained from the Affymetrix platform were normalized using the RMA method in Affymetrix Expression Console software (Affymetrix, CA) (Irizarry et al., 2003). As described by Lyons et al. (2017), gene modules of specific immune cell expression profiles were constructed for the purpose of immune-cell-specific profiling of cancer. With these specific immune modules, we conducted gene-set enrichment analyses using the online java module of GSEA software version 3.0 (Subramanian et al., 2005). A network-based analysis of gene enrichment was performed with Cytoscape software version 3.6.0 (Cline et al., 2007). Differentially expressed genes were identified with a rank products analysis. A functional analysis of genes that were upregulated in the vaccinated group was performed using the Gene Ontology biological process database and the DAVID application from the NIH website (Huang et al., 2009). Bioinformatic analysis was performed in R software version 3.4.1; the R-package made4 was used to construct an expression heatmap using Euclidean distances and the Ward method (Culhane et al., 2005).

### qRT-PCR

Total RNA was isolated using TRIzol Reagent (Life Technologies) and reverse transcription was performed using MultiScribe Reverse Transcriptase (Applied Biosystems) according to the manufacturer’s instructions. Quantitative PCR (qPCR) was performed in duplicate using SYBR Green PCR Master Mix (Applied Biosystems) on an Agilent Technologies Stratagene MX3005p apparatus. The expression levels of genes were normalized to that of glyceraldehyde-3-phosphate dehydrogenase (GAPDH). The sequences of the primers used are: for CXCL9: Fw: CCATGAAGTCCGCTGTTCTT, Rv: TGAGGGATTTGTAGTGGATCG, for CXCL10: Fw: ATCAGCACCATGAACCCAAG, Rv: TTCCCTATGGCCCTCATTCT, for CXCL13: Fw: ATGAGGCTCAGCACAGCA, Rv: ATGGGCTTCCAGAATACCG.

### Immunofluorescence Microscopy

Cells were cultured on glass cover slips (Kniffel Glass) coated with Geltrex (Thermo Fisher). They were fixed with 4% paraformaldehyde and permeabilized with 0.25% Triton 100X; nonspecific sites were blocked with 0.1% PBS Tween+10% FBS. For immunostaining we used the primary antibodies α-Oct4 (Santa Cruz, dilution 1:300) and α-Nanog (Cell Signaling, dilution 1:100) and the secondary antibodies goat anti-mouse Cy3 (Abcam) and goat anti-rabbit Alexa488 (Abcam). Cell nuclei were counterstained with DAPI (4’, 6-diamidino-2-phenylindole) and images were acquired using a Nikon Eclipse 90i microscope.

### Immunohistochemistry of teratomas and quantitative monitoring of lung metastasis

The immunohistochemistry (IHC) of teratomas, was characterized using the methods described (Griscelli et al. 2012). To quantify lung metastasis, lungs were resected, included in paraffin and analyzed after hematoxylin-eosin-safran staining for the detection of lungs metastasis areas. Image digititalized on Nikon Eclipse 90i microscope were used for metastatic areas quantification with Image J software and values obtained with formula (metastatic area of lung/ total area of lung)*100% were compared between different groups.

### Quantification and Statistical Analyses

All values are expressed as mean ± s.e. Differences among groups were assessed, as appropriate, using either an unpaired two-tailed Student’s t test or a one-way/two-way analysis of variance (ANOVA) in PRISM GraphPad software or Microsoft Office Excel software. *P < 0.05, **P < 0.01, ***P < 0.001, ****P < 0.0001.

## DISCUSSION

Although the concept that some cancers might originate from embryonic tissues dates back to the 19^th^ century (Brewer et al. 2009), experimental tests using embryonic tissues as cancer vaccines were not attempted until nearly 100 years later. In early studies, mice were immunized with fetal material (Brewer et al. 2009) and antisera were developed against 9-day-old mouse embryos (Stonehill et al. 1970) that led to the rejection of chemically induced tumors in these animals. With further refinement of the concept of cancer stem cells and the availability of murine and human ESCs, the idea of using PSCs as cancer vaccines has reemerged during the last decade. Several groups have suggested using PSCs as a source of tumor-associated antigens (TAAs), which can prime the immune system to target different types of cancer (Li et al., 2009; Yaddanapudi et al., 2012; Kooreman et al. 2018). Initial experiments were based on the observation that cancer cells and embryonic tissues share a number of cellular and molecular properties, as well as a considerable number of TAAs (de Almeida et al., 2014; Zhao et al., 2011). A major breakthrough came with the discovery of techniques for reprogramming somatic cells to pluripotency; this has made it possible to explore the use of iPSCs as sources of TAAs, which was first tested in an autologous setting in mice (Kooreman al 2018). These experiments, combined with previous studies showing that human ESCs could be used as tumor antigens (Li et al. 2009; Dong et al. 2010), opened doors for the possibility of using PSCs in clinical immunotherapy. To date, multiple ESC and iPSC vaccine candidates have been evaluated, but so far investigations have focused only on their ability to hinder tumor growth. There has been no demonstration of their potential anti-metastatic properties, nor any assessment of whether these vaccines could specifically target CSCs. Clarification of this point was essential, especially in light of the fact that the embryonic genes expressed in several types of CSCs, from aggressive tumors such as non-small cell lung cancer (Hassan et al. 2009), ovarian cancer (Riester et al. 2017) and glioblastoma (Ben-Porath et al. 2008), are also found in ESCs or iPSCs.

In this study, we evaluated for the first time the anti-tumoral potential of a treatment comprised of vaccination with PSCs and administration of the histone deacetylase inhibitor (HDACi) valproic acid (VPA), which has multiple immune stimulatory functions and is able to modify the tumor microenvironment. In particular, we observed that VPA had the potential to significantly increase the expression of MHC1 in 4T1 cells and in CSC/MS-derived 4T1 cells. VPA also increased the expression of MHC2 (CD74), chemokine CCL2, and TNFRSF9, a member of the TNF-receptor superfamily that is known to contribute to the clonal expansion, survival, and development of T cells and to regulate CD28 co-stimulation to promote Th1 cell responses. We also determined that VPA promoted the production *in vivo* of multiple chemokines, such as CXCL9, 10, and 13, which enabled significant modifications in the immunosuppressive microenvironment of tumor cells and facilitated the local recruitment of T and B cells within the tumor.

To evaluate candidates for PSC+VPA treatment, we used a model of an aggressive, poorly differentiated triple-negative breast cancer (TNBC): the 4T1 cell line, which is derived from a spontaneously arising BALB/c mammary tumor (Tao et al. 2008; Aslakson et al. 1992). In this model, transplantation of as few as 5×10^4^ tumor cells into the mammary fat pad of mice results in rapid tumor growth, with swift metastatic spread occurring via a hematogenous route mainly to the lungs and brain; in this, it closely resembles human metastatic breast cancer (Morecki et al. 1998, Wagenblast et al. 2015). Our approach therefore differs from that used by Kooreman et al. (2018), who evaluated the anti-tumor potential of autologous FVB-derived murine iPSCs, combined with a CpG ODN 1826 adjuvant, using a DB7 breast tumor model. That particular cell line has a very low metastatic potential, which does not enable evaluation of the anti-metastatic properties of autologous iPSC-derived vaccines. By using the aggressive 4T1 cell line, we are able to show for the first time not only the tumor-protective abilities of PSCs but also their significant anti-metastatic potential.

In this work, we also evaluated the possibility of generating anti-cancer immunity not just towards “bulk” cancer cells but also towards more primitive CSC-like mammosphere cells.

It is well established that CSCs are the cause of resistance to “classical” therapies, and their persistence lies at the origin of relapses in several types of aggressive cancers (Holohan et al. 2013; Diehn et al. 2009; Bao et al. 2006). It is also well known that CSCs promote metastatic dissemination, which can occur very early on in the progression of cancer with the generation of difficult-to-detect metastatic cells which remain in dormancy (Plaks et al. 2015). Furthermore, it has also been reported that a subpopulation of CSCs undergoes the epithelial-mesenchymal transition (EMT; Bronsert et al. 2014) and acquires a migratory phenotype, which also confers resistance to anti-proliferative drugs (Creighton et al. 2009). Importantly, CSCs that are resistant to conventional therapies (Shafee et al. 2008; Yamauchi et al. 2008) cannot be eradicated by immunotherapy vaccine strategies if such approaches do not introduce to the cancer-bearing host a combination of TAAs that are expressed by the tumor. Thus far, most attempts to develop a cancer vaccine have used TAAs isolated from differentiated cancer cells rather than from CSCs, which will limit the long-term effectiveness of these techniques.

Prior to our study, there had been no evidence presented as to whether immunotherapy vaccine strategies that use PSCs as the source of TAAs have the ability to specifically target CSCs and/or the CSC niche in TNBC. The 4T1 breast cancer model is well suited for addressing this question as it is known to hijack some of the normal stem cell pathways to increase cellular plasticity and stemness (Wagenblast et al. 2015) and thus cause lethal diseases (Sally et al. 2007). To investigate this question, we developed a protocol in which we developed 4T1 cells into CSC-like mammospheres using an EMT-like process, expressing ALDH1 (Ginestier et al. 2007) and exhibiting CSC-like characteristics. Indeed, our 4T1-derived CSCs/MSs had higher colony formation ability and MS formation efficiency than their 4T1 parent cells, and expressed embryonic stem cell-associated factors such as NANOG, which is associated with a poor prognosis in breast cancer patients (Nagata et al. 2014), and SOX2, which is correlated with increased dissemination ability of breast cancer (Liu et al. 2017). In addition, the MSs we generated had higher tumorigenicity *in vivo* than the bulk tumor cells, which suggested that the cell populations contained in the MSs were enriched with more-aggressive breast CSCs.

Our combinatory regimen PSC vaccine with VPA as adjunct had a substantially enhanced anti-tumor effect compared to the vaccine-only treatment, and caused a highly significant reduction in lung metastases (12-fold reduction compared to mice treated with the vaccine without VPA, Figure 4E). These results indicate that VPA significantly modified the immunosuppressive microenvironment within the primary tumor and reduced the number of cancer cells with an EMT/CSC phenotype that were able to migrate to the secondary organs. This anti-metastatic effect was associated with an increase in CD4^+^ and CD8^+^ T cells in tumors and spleens, as well as a significant decrease of PD1^+^ in CD4^+^ and CD8^+^ T cells in the secondary lymphoid organs, which was detected only in the group of mice that received VPA. One of the results of our work that is, to our knowledge, unprecedented is the demonstration that the combination of PSC and HDACi therapy enables the induction of effective anti-tumor immunity against 4T1-derived CSCs/MSs. Indeed, this combinatory regimen was highly effective in preventing the establishment of 4T1-derived CSCs/MSs, and this result was significantly correlated with an increase in CD4^+^ and CD8^+^ T cells and IFN-γ-secreting cytotoxic T lymphocytes, along with the attenuation of Gr1^+^CD11b^+^ MDSCs within the tumor. Our study also revealed that hESCs were more effective than mESCs in generating immunity to TNBC in naïve mice, probably due to increased heterogeneity in TAA expression in hESCs compared to mESCs and/or to improved cross-presentation of human material on murine MHC or H-2 molecules, which then improved activation of host CD4^+^ and CD8^+^ T cells. To address this, further research is needed to investigate the mechanism of antigen cross-presentation from hESCs by the dedicated host antigen-presenting cells, such as dendritic cells, as well as the processes behind the subsequent cross-priming of antigen-specific T cells.

Because the use of embryonic materials to create vaccines could incite ethical and political controversies, we also evaluated iPSCs as a source of TAAs, and used these cells to immunize naïve mice against a lethal dose of 4T1 breast carcinoma cells. For this purpose, we evaluated two sources of fibroblast-derived iPSCs that were generated from the BALB/c and C57BL/6 strains of mice, respectively; this enabled us to compare anti-tumor immunity generated in BALB/c mice under autologous versus allogeneic conditions. Regardless of the strain of origin, vaccinated mice had significantly smaller tumors compared to unvaccinated controls, and the inclusion of VPA treatment increased the anti-tumoral response considerably. We also found a significant improvement in the survival rate due to vaccination, and this was more pronounced in mice that had been primed with allogeneic material (versus autologous) combined with VPA. Another major finding was that vaccination with miPSCs was able to generate an effective memory immune response against breast 4T1 breast carcinoma. Mice that were challenged with 5×10^4^ 4T1-GFP-Luc cells 30 or 120 days after the end of a six-dose vaccination series were more able to reject CSC/MS-derived 4T1 cells than unvaccinated mice; this result indicated that our vaccination protocol had established long-term immune memory and led to the activation of anti-tumor immunity, mainly against CSCs.

Interestingly, allogeneic iPSCs appeared to be more effective than autologous iPSCs in preventing the establishment of 4T1 carcinoma cells. This could represent a considerable advantage in efforts to create a clinical vaccine to prevent and treat TNBC. Indeed, allogeneic iPSCs may facilitate the development of “off-the-shelf” material that could be used for curative approaches for which time is of the essence, such as in the treatment of relapses. Autologous-derived iPSCs would likely be useful only in prophylactic settings, as the length of time needed to produce bona fide iPSCs (several months, corresponding to several passages in culture) would render them highly impractical for treatment. Moreover, their use in a clinical setting would also require the use of multiple quality control assays to ensure their purity and safety, which would create further delays.

In our study, we did not observe any significant autoimmunity: immunized mice were generally healthy and presented no clinical evidence of autoimmune diseases. The animals’ weight, hair, and musculature were normal. However, more follow-up is needed before iPSC-based cancer vaccines move into clinical testing.

Taken together, our data show the feasibility of creating anti-cancer immunity through an approach that combines an HDACi and a PSC-based vaccine. This technique can be easily applied to patients and, because it targets CSCs, significantly decreases the occurrence of metastasis. This combinatory regimen could increase the survival of certain patients with CSC-phenotype tumors, particularly given that for solid tumors, invasion and metastasis account for more than 90% of mortality (Bronsert et al. 2014; Sleeman, et al. 2010; Lazebnik et al. 2010).

In addition, our data indicate that this regimen of PSC vaccine-and-HDACi led to significant modifications in the tumor microenvironment, by recruiting T and B cells and by modifying certain stromal cellular components. That can be the case of myeloid-derived suppressor cells in the primary tumor and distant organs, which have been shown to regulate tumor plasticity by inducing an EMT/CSC phenotype, thus facilitating the dissemination of tumor cells from the primary site (Ouzounova et al. 2017). Likewise, these effects may hinder the activity of tumor-associated macrophages, which are able to produce pro-invasive cytokines that not only affect invasion directly but also sustain the cancer-associated mesenchymal phenotype (Noy et al. 2016). These beneficial properties make this combinatory regimen a potentially powerful option for a new concept of immunotherapy that could be deployed shortly after conventional primary treatment of cancer or in combination with conventional therapies or “checkpoint inhibitors”, which are currently under intensive investigation.

## ACKNOWLEDGMENTS

We sincerely thank Olivia Bawa, Mathis Soubeyrand, Olivier Feraud, and Dominique Divers for technical assistance; Benoit Peuteman for animal care; Paule Opolon for anatomopathological analysis; and Jerome Artus for help in deriving miPSCs. This work was performed with grants from ANR Programme d’Investissements d’Avenir of the INGESTEM National Infrastructure Program (ANR-11-INBS-0009-INGESTEM), Inserm, University Paris Sud, and Vaincre le Cancer NRB.

## AUTHOR CONTRIBUTIONS

Study concept and financing: F.G., A.B.G., A.T. Direction of the project: F.G., A.B.G, A.T. Generation, production, and characterization of miPSCs: M.H.K. Molecular analysis: C.D. Mouse experiments: A.A, M.H.K, F.G. Cell profiles: M.H.K, A.A, M-G.G.H. Flow cytometry: A.A, M.H.K, D.C, M-G.G.H. Microarray: C.D. Writing of the article: F.G, A.B.G, A.T., M.H.K. qRT-PCR: A.A.

## FIGURE LEGENDS

**Supplementary figure 1. Characterization of D3 murine embryonic stem cells.**

(A) Morphology of D3 cells under 40X magnification, typical of ESC colonies.

(B) Expression of the key pluripotency markers NANOG and OCT4 was visualized using immunofluorescence.

(C) Murine ESCs exhibited typical markers of pluripotency, such as the expression of SSEA1, as quantified by flow cytometry.

(D) The ability to differentiate into the three germ layers was confirmed by teratoma formation assays, which revealed differentiation into ectodermal, endodermal, and mesodermal tissues.

**Supplementary figure 2. VPA treatment increased MHC I expression**

(A) Increase in MHC I expression on the surface of 4T1 cells as a result of treatment with different doses of VPA (0 mM, 0.2 mM, and 2 mM), as revealed by flow cytometry

(B) Effect of treatment with 2 mM VPA on the expression of MHC I in adherent 4T1 cells and 4T1-derived MSs.

**Supplementary figure 3. VPA-only treatment did not hinder *in vivo* tumor growth.**

(A) Tumor growth (mm^3^) in mice treated with VPA compared to control mice (treated with PBS). There was no significant differences between groups.

(B) IVIS imaging of VPA-treated and untreated mice.

(C) Lung metastases in VPA-treated and untreated mice were quantified using bioluminescence imaging. Regions of interest (ROI) for pulmonary metastases in these two groups were calculated by Living Image Software.

(D) Effects of VPA treatment on the frequencies of CD4^+^ cells, CD8^+^ cells, Tregs, and MDSCs compared to controls, as quantified by flow cytometry.

**Supplementary figure 4. Impact of hESC vaccination with CpG adjunct on tumor formation and lung metastasis**

(A) Illustration of the protocol for hESC vaccination and CpG administration: CpG was mixed with 2×10^6^ irradiated hESC cells and injected in mice twice, with a one-week interval between injections. 4T1-GFP-Luc tumor cells were then implanted in the mammary fat pad of mice.

(B) IVIS imaging of mice vaccinated with hESCs+CpG compared to the control group

(C) Tumor volume in mice treated with hESCs+CpG compared to controls.

(D) Tumor weight in mice treated with hESC+CpG compared to controls.

(E) Quantification (total flux) of luciferase by bioluminescence imaging of the lungs from mice vaccinated with hESC+CpG compared to controls

(F) Correlation between tumor weight and metastatic spread in the lungs.

**Supplementary figure 5. Generation of mammospheres from the 4T1 cell line.**

(A) Images of the morphology of adherent 4T1 cells and 4T1-derived mammospheres in 3D culture conditions, cultured with or without TGFβ + TNFα (magnification x20).

(B) Quantification of ALDH1 activity by flow cytometry in adherent 4T1 cells and in 4T1-derived mammospheres cultured with or without TGFβ + TNFα.

(C) Percentage of ALDH1^+^ cells among adherent 4T1 cells and 4T1-derived mammospheres cultured with or without TGFβ + TNFα. Results are shown from three independent experiments.

(D) Mammosphere-formation efficiency (calculated by Cell Selector Software) indicates the number of mammospheres of different sizes obtained with or without TGFβ +TNFα treatment.

(E) Quantification of mRNA expression of NANOG, OCT4, and SOX2 by qRT-PCR in adherent 4T1 cells and in 4T1-derived mammospheres cultured with or without TGFβ + TNFα.

**Supplementary figure 6. Evaluation of the anti-tumor and anti-metastatic potential of human or murine ESCs in combination with VPA in mice challenged with 4T1-derived mammospheres grown in 3D culture conditions with TGFβ + TNFα.**

(A) Comparison of tumor growth in mice injected with 2.5×10^4^ adherent 4T1 cells or 4T1-derived mammospheres produced using TGFβ + TNFα.

(B) Tumor volumes in mice treated with hESCs+VPA or mESCs+VPA compared to untreated controls. All mice were challenged with 2.5×10^4^ 4T1-derived mammospheres that were produced using TGFβ + TNFα. At day 23 post-challenge, treated mice had significantly smaller tumors (p<0.001).

(C) Bioluminescence imaging of mice vaccinated with hESCs+VPA compared to unvaccinated control mice.

(D) Quantification of lung metastatic areas at day 23.

(E) Direct bioluminescence imaging of surgically removed lungs showed a reduction in or the absence of metastastic load in the lungs of mice treated with hESCs+VPA compared to untreated controls.

(F) Hematoxylin and eosin staining revealed lung metastases in mice treated with human or murine ESCs+VPA and those in untreated controls.

**Supplementary figure 7. Characterization of murine induced pluripotent stem cells derived from BALB/c fibroblasts**

(A) Morphological view under the microscope of miPSCs expanded on mouse embryonic fibroblasts.

(B) Expression of the key pluripotency markers NANOG and OCT4 as revealed by immunofluorescence; DAPI was used as counterstain.

(C) Expression of SSEA1 in cell membranes as quantified by flow cytometry.

(D) Teratoma formation assays, showing differentiation into ectodermal, endodermal, and mesodermal tissues.

**Supplementary figure 8. Effective memory immune response following vaccination with miPSCs**

(A) Experimental protocol to evaluate *in vivo* immune memory generated by vaccination: BALB/c mice were injected subcutaneously six times with 2×10^6^ miPSCs (15Gy irradiated) in the right flank. Tumor challenge occurred 30 days after the final vaccine dose was given; mice were injected with 2.5×10^4^ 4T1-GFP-Luc cells and VPA was added in their drinking water.

(B) At day 26 post-challenge, breast tumors were significantly smaller in mice that had undergone the six-month vaccination protocol compared to unvaccinated mice (n=5 per group).

(C) Survival curves of miPSC+VPA-treated mice or PBS-treated controls.

**Supplementary figure 9. Immune cell profiling of the spleens of mice treated with miPSCs+VPA**

(A, B) Frequency of CD4^+^ and CD8^+^ T cells in the spleens of mice treated with miPSCs+VPA compared with untreated controls, as measured by flow cytometry.

(C) Frequency of PD1 in the cell membranes of CD4^+^ T cells, as measured by flow cytometry.

(D) Frequency of PD1^+^ T-effector memory (CD44^+^CD62L^-^CCR7^low^CD127^low^) cells in the spleen, as measured by flow cytometry.

**Supplementary figure 10.**

(A) Experimental protocol to evaluate *in vivo* immune memory generated by vaccination: BALB/c mice were injected subcutaneously six times with 2×10^6^ miPSCs (15Gy irradiated) in the right flank. Tumor challenge occurred 120 days after the final vaccine dose was given; mice were injected with 2.5×10^4^ 4T1-GFP-Luc-derived mammospheres and VPA was added in their drinking water.

(B) At day 28 post-challenge, breast tumors were significantly smaller in mice that had undergone the six-month vaccination protocol compared to unvaccinated mice (n=6 per group).

(G) Bioluminescence imaging of tumors from mice treated with miPSCs+VPA compared to untreated controls.

(H) Bioluminescence imaging of lungs isolated from vaccinated and control mice 28 days after challenge.

(I) A significant correlation was found between tumor burden and metastatic spread at day 28.

**Supplementary figure 11. iPSC+VPA treatment induced the recruitment of immune cells by chemokines.**

Results of qRT-PCR showing strong upregulation of CXCL9, CXCL10, and CXCL13 mRNA as a result of miPSC+VPA treatment.

**Table 1. Differentially expressed immune genes that were upregulated in 4T1 cells by treatment with valproic acid**. The list highlights immune-associated genes that were found to be significantly overexpressed in the treatment group using the Significance Analysis of Microarray algorithm with a false discovery rate of less than 5%. Columns include: gene symbol, gene description, gene ID number, and the fold-change in expression between the VPA-treated group and untreated controls.

**Table 2. Genes that were differentially expressed between vaccinated and control mice that developed 4T1-derived tumors**: Differentially expressed genes are listed with their gene symbol, description, the fold-change in expression found in transcriptome experiments (Vaccine/Control), and the p-value of the comparison.

