## Supplementary Tables for "Pharmacologically modified pluripotent stem cell-based cancer vaccines with anti-metastatic potential"

Table 1

| Gene symbol | Description | FC vaccin/CT 4T1 | Ranking p-values |
|---|---|---|---|
| Cxcl13 | chemokine (C-X-C motif) ligand 13 | 8,877 | 5.1787803E-5 |
| Gm17482 | predicted gene, 17482 [Source:MGI Symbol;Acc:MGI:4937116] | 3,117 | 1.3509862E-6 |
| Mir675 | microRNA 675; H19, imprinted maternally expressed transcript | 2,667 | 1.8913806E-5 |
| Clec4a1 | C-type lectin domain family 4, member a1 | 2,235 | 1.6932361E-4 |
| Fxyd1 | FXYD domain-containing ion transport regulator 1 | 2,042 | 2.6119065E-5 |
| Il20rb | interleukin 20 receptor beta | 2,014 | 3.3324326E-5 |
| Sh2d4a | SH2 domain containing 4A | 1,945 | 5.439971E-4 |
| Egln3 | EGL nine homolog 3 | 1,939 | 4.6158695E-4 |
| Serpinb2 | serine (or cysteine) peptidase inhibitor, clade B, member 2 | 1,886 | 3.449518E-4 |
| Clec12a | C-type lectin domain family 12, member a | 1,840 | 0.0011415834 |
| Lmo7 | LIM domain only 7 | 1,828 | 1.6707196E-4 |
| Vps33b | vacuolar protein sorting 33B (yeast) | 1,828 | 3.62965E-4 |
| Pmp22 | peripheral myelin protein 22 | 1,815 | 1.1618481E-4 |
| Ghr | growth hormone receptor | 1,809 | 7.3583715E-4 |
| Ly96 | lymphocyte antigen 96 | 1,759 | 7.488967E-4 |
| Cldn9 | claudin 9 | 1,747 | 2.193101E-4 |
| 9230104L09Rik | RIKEN cDNA 9230104L09 gene | 1,741 | 2.3011799E-4 |
| Ccl22 | chemokine (C-C motif) ligand 22 | 1,729 | 4.0664684E-4 |
| Ttll1 | tubulin tyrosine ligase-like 1 | 1,717 | 8.475187E-4 |
| Naf1 | nuclear assembly factor 1 homolog | 1,711 | 6.4351974E-4 |
| Enpp2 | ectonucleotide pyrophosphatase/phosphodiesterase 2 | 1,699 | 6.5387733E-4 |
| Tc2n | tandem C2 domains, nuclear | 1,699 | 0.0012348014 |
| Slc6a8 | solute carrier family 6 (neurotransmitter transporter, creatine), r | 1,693 | 8.2500227E-4 |
| Shroom3 | shroom family member 3 | 1,688 | 8.290552E-4 |
| Zfp595 | zinc finger protein 595 | 1,688 | 6.4351974E-4 |
| Sod3 | superoxide dismutase 3, extracellular | 1,676 | 4.3952087E-4 |
| Cd46 | CD46 antigen, complement regulatory protein | 1,670 | 6.642349E-4 |
| Cd72 | CD72 antigen | 1,670 | 0.0017567324 |
| Pid1 | phosphotyrosine interaction domain containing 1 | 1,670 | 0.0011983248 |
| Psmg1 | proteasome (prosome, macropain) assembly chaperone 1 | 1,664 | 0.0011478879 |
| Rnf223 | ring finger 223 | 1,664 | 6.421688E-4 |
| Cd300e | CD300e antigen | 1,641 | 0.0011577952 |
| Csf2ra | colony stimulating factor 2 receptor, alpha | 1,641 | 0.002091777 |
| Hnmt | histamine N-methyltransferase | 1,641 | 0.0011888678 |
| Cpa3 | carboxypeptidase A3, mast cell | 1,630 | 0.0023011798 |
| Olfml2b | olfactomedin-like 2B | 1,625 | 0.0013100064 |
| Rassf9 | Ras association (RalGDS/AF-6) domain family (N-terminal) memb | 1,625 | 0.001398721 |
| Bin2 | bridging integrator 2 | 1,619 | 0.0028267135 |
| Ms4a14 | PREDICTED: membrane-spanning 4-domains, subfamily A, mem | 1,619 | 0.0020715122 |
| Olfr384 | olfactory receptor 384 | 1,613 | 7.7321444E-4 |
| Hormad2 | HORMA domain containing 2 | 1,608 | 0.0018985859 |
| Gfra2 | glial cell line derived neurotrophic factor family receptor alpha 2 | 1,597 | 0.0012388544 |
| Blk | B lymphoid kinase | 1,586 | 9.051608E-4 |
| Ms4a7 | membrane-spanning 4-domains, subfamily A, member 7 | 1,569 | 0.0021237503 |
| Smpd4 | sphingomyelin phosphodiesterase 4 | 1,564 | 0.0023291004 |
| Ccr2 | chemokine (C-C motif) receptor 2 | 1,553 | 0.0021359092 |
| Hhip | Hedgehog-interacting protein | 1,553 | 0.0018076196 |
| Ctse | cathepsin E | 1,548 | 0.0041119517 |
| Gm2573 | PREDICTED: predicted gene 2573 (Gm2573) | 1,542 | 0.003948032 |

| | | | |
|---|---|---|---|
| Rasl12 | RAS-like, family 12 | 1,542 | 0.0016527065 |
| Smtnl1 | smoothelin-like 1 | 1,542 | 0.0025668738 |
| Gm5595 | predicted gene 5595 | 1,537 | 0.002425921 |
| Pbx4 | pre B cell leukemia homeobox 4 | 1,537 | 0.0018666127 |
| Etv3 | ets variant 3 | 1,532 | 0.002837071 |
| Fam49a | family with sequence similarity 49, member A | 1,532 | 0.0028897594 |
| Msmp | microseminoprotein, prostate associated | 1,532 | 0.0013523372 |
| Cdon | cell adhesion molecule-related/down-regulated by oncogenes | 1,526 | 0.0024592453 |
| Atp13a5 | ATPase type 13A5 | 1,521 | 0.002064307 |
| Inmt | indolethylamine N-methyltransferase | 1,521 | 0.0027361973 |
| Gm1818 | predicted gene 1818 [Source:MGI Symbol;Acc:MGI:3037676] | 1,516 | 0.0032720887 |
| Nupl2 | nucleoporin like 2 | 1,516 | 0.004172296 |
| Pgr15l | G protein-coupled receptor 15-like | 1,516 | 0.0024200666 |
| Trp53i11 | transformation related protein 53 inducible protein 11 | 1,516 | 0.004660002 |
| Aldh2 | aldehyde dehydrogenase 2, mitochondrial | 1,510 | 0.0034625777 |
| Txnrd3 | thioredoxin reductase 3 | 1,510 | 0.0027334956 |
| Aim2 | absent in melanoma 2 | 1,510 | 0.002627218 |
| Fhit | fragile histidine triad gene | 1,510 | 0.0038422048 |
| Entpd5 | ectonucleoside triphosphate diphosphohydrolase 5 | 1,505 | 0.0027798794 |
| Gjb4 | gap junction protein, beta 4 | 1,505 | 0.0046388363 |
| Ndufa4l2 | NADH dehydrogenase (ubiquinone) 1 alpha subcomplex, 4-like 2 | 1,505 | 0.0017959111 |
| Olfr822 | olfactory receptor 822 | 1,505 | 0.0020602539 |
| Pou4f1 | POU domain, class 4, transcription factor 1 | 1,500 | 0.004528956 |
| Syt13 | synaptotagmin XIII | 1,500 | 0.0029388454 |
| Dpep1 | dipeptidase 1 (renal) | 1,495 | 0.0030883546 |
| Olfr1258 | olfactory receptor 1258 | 1,495 | 0.0026402774 |
| Fbln5 | fibulin 5 | 1,490 | 0.0046690083 |
| Krt14 | keratin 14 | 1,490 | 0.002484914 |
| Olfr452 | olfactory receptor 452 | 1,490 | 0.003948032 |
| Tuft1 | tuftelin 1 | 1,490 | 0.0048991265 |
| Ncoa7 | nuclear receptor coactivator 7 | 1,485 | 0.003563001 |
| Nipal3 | NIPA-like domain containing 3 | 1,485 | 0.004543367 |
| Mest | mesoderm specific transcript | 1,479 | 0.0041196076 |
| Mtfr2 | mitochondrial fission regulator 2 | 1,474 | 0.0044375393 |
| Ssx2ip | synovial sarcoma, X breakpoint 2 interacting protein | 1,474 | 0.0041115014 |
| Stard3 | START domain containing 3 | 1,474 | 0.0044524004 |
| Wasf1 | WAS protein family, member 1 | 1,474 | 0.0032923534 |
| Ica1 | islet cell autoantigen 1 | 1,469 | 0.0048032063 |
| Nmrk1 | nicotinamide riboside kinase 1 | 1,469 | 0.0043871026 |
| Sord | sorbitol dehydrogenase | 1,469 | 0.0047469153 |
| Kdm1b | lysine (K)-specific demethylase 1B | 1,464 | 0.0040673693 |
| Slx4ip | SLX4 interacting protein | 1,464 | 0.0049171397 |
| Klra17 | killer cell lectin-like receptor, subfamily A, member 17 | 1,459 | 0.0037521392 |
| Smr2 | submaxillary gland androgen regulated protein 2 | 1,459 | 0.0045280554 |
| Pigw | phosphatidylinositol glycan anchor biosynthesis, class W | 1,454 | 0.0044258307 |
| Ano2 | anoctamin 2 | 1,444 | 0.0044839233 |
| Klk1b11 | kallikrein 1-related peptidase b11 | 1,444 | 0.004620823 |
| Ppp1r3f | protein phosphatase 1, regulatory (inhibitor) subunit 3F | 1,444 | 0.0031455462 |
| 1700011H14Rik | RIKEN cDNA 1700011H14 gene | 1,439 | 0.004207872 |

Table 2

| Gene symbol | Description | Gene ID | Fold change (Valproic acid /control) |
|---|---|---|---|
| CD74 | CD74 molecule | 972 | 2.756137 |
| TNFRSF9 | TNF receptor superfamily member 9 | 3604 | 2.152303 |
| CCL2 | C-C motif chemokine ligand 2 | 6347 | 2.0778043 |
| CASP4 | caspase 4 | 837 | 1.812713 |
| AREG | amphiregulin | 374 | 1.7866501 |
| IFIT2 | interferon induced protein with tetratricopeptide repeats 2 | 3433 | 1.7681799 |
| SERPINB2 | serpin family B member 2 | 5055 | 1.7294099 |
| LY6E | lymphocyte antigen 6 family member E | 4061 | 1.7189492 |
| PLK2 | polo like kinase 2 | 10769 | 1.6721709 |
| EGR1 | early growth response 1 | 1958 | 1.6136316 |
| DRAM1 | DNA damage regulated autophagy modulator 1 | 55332 | 1.6093787 |
| KLF9 | Kruppel like factor 9 | 687 | 1.6086748 |
| IFIH1 | interferon induced with helicase C domain 1 | 64135 | 1.5915328 |
| UBE2L6 | ubiquitin conjugating enzyme E2 L6 | 9246 | 1.559597 |
| IL15RA | interleukin 15 receptor subunit alpha | 3601 | 1.5306206 |
| CCRL2 | C-C motif chemokine receptor like 2 | 9034 | 1.524512 |
| STAT1 | signal transducer and activator of transcription 1 | 6772 | 1.4973801 |
| NLRC5 | NLR family CARD domain containing 5 | 84166 | 1.4913481 |
| ISOC1 | isochorismatase domain containing 1 | 51015 | 1.4899585 |
| BST2 | bone marrow stromal cell antigen 2 | 684 | 1.4610225 |
| IL15 | interleukin 15 | 3600 | 1.4604695 |
| SAMD9L | sterile alpha motif domain containing 9 like | 219285 | 1.448133 |
| HERC6 | HECT and RLD domain containing E3 ubiquitin protein ligase family member 6 | 55008 | 1.4128106 |
| DHX58 | DExH-box helicase 58 | 79132 | 1.4095724 |
| CSF1 | colony stimulating factor 1 | 1435 | 1.401427 |
| SOD2 | superoxide dismutase 2 | 6648 | 1.3938667 |
| PSMB9 | proteasome subunit beta 9 | 5698 | 1.3776621 |
| SLC2A6 | solute carrier family 2 member 6 | 11182 | 1.3619148 |
| RNF213 | ring finger protein 213 | 57674 | 1.3537916 |
| IL7 | interleukin 7 | 3574 | 1.3391039 |
| TUBB2A | tubulin beta 2A class IIa | 7280 | 1.331755 |
| TDRD7 | tudor domain containing 7 | 23424 | 1.3295585 |
| ATP2B1 | ATPase plasma membrane Ca2+ transporting 1 | 490 | 1.3081365 |
| CCL20 | C-C motif chemokine ligand 20 | 6364 | 1.307062 |
| CD274 | CD274 molecule | 29126 | 1.2911943 |
| CEBPD | CCAAT/enhancer binding protein delta | 1052 | 1.2831194 |
| ZBTB10 | zinc finger and BTB domain containing 10 | 65986 | 1.2745908 |
| IRS2 | insulin receptor substrate 2 | 8660 | 1.2700346 |
| NCOA7 | nuclear receptor coactivator 7 | 135112 | 1.2286383 |
| REL | REL proto-oncogene, NF-kB subunit | 5966 | 1.2227736 |
| CCL7 | C-C motif chemokine ligand 7 | 6354 | 1.1728142 |
| IL10RA | interleukin 10 receptor subunit alpha | 3587 | 1.1703566 |
| SP110 | SP110 nuclear body protein | 3431 | 1.1379845 |
| MSC | musculin | 9242 | 1.1052907 |
